# Diversity and seasonality of ectoparasite burden on two species of Madagascar fruit bat, *Eidolon dupreanum* and *Rousettus madagascariensis*

**DOI:** 10.1101/2025.01.20.633693

**Authors:** Angelo F. Andrianiaina, Santino Andry, Gwenddolen Kettenburg, Hafaliana Christian Ranaivoson, Vincent Lacoste, Philippe Dussart, Jean-Michel Heraud, Theresa M. Laverty, Sarah Guth, Katherine I. Young, Aristide Andrianarimisa, Cara E. Brook

**Affiliations:** Department of Zoology and Animal Biodiversity, University of Antananarivo, Madagascar; Department of Entomology, University of Antananarivo, Madagascar; Department of Ecology and Evolution, University of Chicago, IL, United States; Virology Unit, Institut Pasteur de Madagascar, Antananarivo, Madagascar; Department of Fish, Wildlife and Conservation Ecology, New Mexico State University, Las Cruces, NM, United States; Department of Biology, Skyline College, San Bruno, CA, United States; Department of Biological Sciences, El Paso, TX, United States

**Keywords:** Bat fly, bat ectoparasite, DNA barcoding, *Eidolon dupreanum*, Madagascar, Nycteribiidae, Pteropodidae, *Rousettus madagascariensis*, Streblidae

## Abstract

**Background:** Bats are important reservoir hosts for a variety of microparasites, some of which are transmitted by ectoparasite vectors that include mites, fleas, lice, ticks, and bat flies (families Nycteribiidae and Streblidae). All of these ectoparasite taxa are known to parasitize two endemic fruit bats of Madagascar, *Eidolon dupreanum* and *Rousettus madagascariensis.* We aimed to describe the diversity of ectoparasite infestation for both bat species through morphological observation and DNA barcoding and elucidate ecological and climatic correlates of seasonal nycteribiid parasitism of these hosts.

**Methods:** Live *E. dupreanum* and *R. madagascariensis* fruit bats were captured monthly in northern and central-eastern Madagascar from 2013-2020. Ectoparasites on all captured bats were counted and identified in the field, then collected into ethanol. Field identification of a subset of samples were confirmed via microscopy and DNA barcoding of the cytochrome C oxidase subunit 1 (COI) and 18S genes. The seasonal abundance of nycteribiid bat flies on both host bats was analyzed using generalized additive models, and the role of climate in driving this seasonality was assessed via cross-correlation analysis combined with generalized linear models. Phylogenetic trees were generated to compare COIand 18S sequences of Madagascar nycteribiid and streblid bat flies with available reference sequences from GenBank.

**Results:** Ectoparasites corresponding to four broad taxa (mites, ticks, fleas, and bat flies) were recovered from 628 of 873 *E. dupreanum* and 831 of 862 *R. madagascariensis*. *E. dupreanum* were most commonly parasitized by *Cyclopodia dubia* nycteribiids and *R. madagascariensis* by *Eucampsipoda madagascariensis* nycteribiids or *Megastrebla wenzeli* streblids. We observed significant seasonality in nycteribiid abundance on both bat hosts, which varied by bat sex and was positively correlated with lagged temperature, precipitation, and humidity variables. Barcoding sequences recovered for all three bat fly species grouped with previously reported sequences, confirming morphological species identification. Our study contributes the first DNA barcodes of any kind reported for *M. wenzeli* and the first 18S barcodes for *C. dubia*.

**Conclusion:** This study explores the diversity and abundance of ectoparasite burdens in two Malagasy fruit bat species, highlighting the importance of seasonal ecology and the influence of climate variables on parasitism, which correlates with resource availability.

## BACKGROUND

Bats (order: Chiroptera) are reservoir hosts for several highly virulent zoonotic viruses (1), which they appear to host without experiencing clinical disease (2). Bats also host a variety of non-viral pathogens, including protozoa (3), bacteria (4), fungi (5,6), and helminths (7), some of which are known to cause significant pathology (5,6,8). In addition, bats can be parasitized by a diversity of hematophagous ectoparasites—including mites, fleas, lice, ticks, and bat flies—which can function as microparasite vectors (9,10). While the majority of bat viruses described to date are directly transmitted via contact with bat excreta (feces, urine, saliva), bat ectoparasites have been reported as potential vectors of *Polychromophilis* spp. (3,11) and *Trypanosoma* spp. (12–15) protozoa, as well as *Bartonella* spp. (4,16–24), *Rickettsia* spp. (16,25,26) and *Borrelia* spp. bacteria (25,25–28). Bat flies (Order: Diptera; Superfamily: Hippoboscoidea) are the most widely recognized bat ectoparasites; bat flies are obligate pupiparous ectoparasites that comprise two families: the monophyletic and wingless Nycteribiidae, which are occasionally found in the New World but most commonly identified on Old World bats, and the paraphyletic, winged Streblidae, for which disparate New World and Old World clades are recognized, with higher diversity in the New World (29,30).

Madagascar, an isolated island off the southeastern coast of Africa, is home to 49 bat species, including 38 endemics (31). Intensive biosurveillance of the island’s bats over the past two decades has led to the discovery and characterization of numerous bat-borne viruses (32–42), protozoa (3,43), bacteria (4,44,45), and helminths (46). Some highly divergent Malagasy bat pathogens reflect the island’s longstanding phylogeographic isolation (34,35,47,48), while others with high identity to African bat pathogens suggest recent cross-continental genetic exchange (34,36,47,48). Parasitism of Malagasy bats by mites, fleas, ticks, and bat flies has been previously described (4,49). Most prior research on Madagascar’s bat flies has focused on elucidating Nycteribiidae diversity (50–53), including through molecular characterization (46,49). At least nine species of nycteribiid bat fly infest a variety of Malagasy bats, including *Cyclopodia dubia* and *Eucampsipoda madagascariensis,* species-specific ectoparasites of, respectively, the endemic Malagasy fruit bats, *Eidolon dupreanum* and *Rousettus madagascariensis* (4,49,54). In addition, distinct *Basilia* sp. nycteribiids have been identified as species-specific ectoparasites of the endemic vesper bats, *Scotophilus robustus*, *S. marovaza*, and *Pipistrellus hesperidus*, as well as the pan-African emballonurid bat, *Taphozous mauritianus* (49,54,55). By contrast, at least three nycteribiid ectoparasites (*Penicillidia* sp., *P. leptothrinax,* and *Nycteribia stylidopsis*) are known to parasitize multiple species of Malagasy bat hosts, including *Myotis goudoti* and at least eight *Miniopterus* spp. (49,54). Only a few early morphological studies describe parasitism of Malagasy bats by Streblidae bat flies: the streblid *Megastrebla wenzeli* has been cited as a species-specific ectoparasite of *R. madagascariensis* (56,57), but to our knowledge, no molecular data for Malagasy streblids have yet been contributed to the literature.

In addition to systematics and taxonomy, several studies have described potential vector roles for Malagasy bat flies. *Bartonella* spp. bacteria have been identified in the Malagasy bat flies, *C. dubia, Basilia* sp., *P. leptothrinax,* and *N. stylidiopsis* (4,58). Our team recovered nested sequences of *Bartonella* spp. in *C. dubia* and their obligate *E. dupreanum* fruit bat hosts; *Bartonella* spp. were absent from *Thaumapsylla* sp. fleas infesting the same bats, suggesting a possible vectorial function for the bat flies (4). Similarly, nested sequences of *Polychromophilus melanipherus* protozoa were detected in *P. leptothrinax* and *N. stylidiopsis* and their obligate bat hosts, *Miniopterus aelleni, M. manavi,* and *M. gleni*, again suggesting a vectorial capacity for the bat flies (3). To our knowledge, experimental confirmation of true vector-microparasite relationships has not yet been carried out for any Malagasy bat fly.

More recent work has provided deeper insights into the ecology of parasite-host relationships for the nycteribiid *E. madagascariensis* and the streblid *M. wenzeli* with their obligate *R. madagascariensis* bat hosts. Prior work in northern Madagascar’s Ankarana National Park showed higher rates of *E. madagascariensis* parasitism of *R. madagascariensis* male vs. female bats and a higher prevalence of parasitism during sampling events carried out in the Malagasy dry (September) vs. wet season (January) (59); the sex ratios of *E. madagascariensis* also skewed towards males (59). No significant differences in parasitism intensity across host sex or age categories or time of sampling were identified for the much less prevalent *M. wenzeli* bat flies (59). Within each sampling season, the same study identified a significant positive correlation between bat body condition index (a proxy for bat health) and the abundance of *E. madagascariensis* (60). Bats with better body conditions were associated with higher abundance bat fly infestations, a result the authors explained could be simply due to due to larger surface area available or improved bat pelage for nycteribiid fixation (60). Field studies in both northern (60) and central Madagascar (61) have shown that fruit bat body conditions improve during Madagascar’s resource-abundant wet season (∼December – April), as compared to the resource-poor dry season. Prior work in northern Madagascar has also documented *R. madagascariensis* consumption of both nycteribiid and streblid bat flies, a habit which provides a likely important protein source to these frugivorous bats in the dry season (62–64).

Here, we aimed to characterize ecological patterns of ectoparasite-host association for two species of cave-dwelling, endemic Malagasy fruit bats, *E. dupreanum* and *R. madagascariensis.* Using data from longitudinally-monitored roost sites in northern and central Madagascar, we sought to quantify seasonal variation in bat fly parasitism for these two bat host species and elucidate a possible role for climate in explaining this variability. Finally, we aimed to expand prior molecular studies of Madagascar bat flies to include *M. wenzeli* streblid parasites of *R. madagascariensis*.

## METHODS

### Bat sampling and ectoparasite collection

Endemic Malagasy fruit bats were captured at roughly six-week intervals at longitudinally-monitored roost sites in central-eastern Madagascar (*Eidolon dupreanum*: Angavokely, Angavobe, Lakato caves; *Rousettus madagascariensis*: Maromizaha cave) and in Ankarana National Park in northern Madagascar between November 2013 – March 2020 in part with ongoing studies characterizing seasonal viral dynamics in these bat populations (Table S1) (32,34–36). Bats were captured using mist nets hung at cave entrances at dusk and dawn. Upon capture, all bats were removed from nets and placed individually in clean, cloth bags for processing (all bags were washed prior to reuse on a new individual). During processing, bats were weighed (in g) using a Pesola scale, and forearm measurements (in mm) were collected with a caliper. All bats were visually examined for ectoparasites, and any observed ectoparasites were removed with foreceps and counted in the field into broad taxonomic categories (ticks, mites, fleas, and bat flies in family Nycteribiidae or Streblidae). Following counting, all ectoparasites collected from a single bat were stored collectively in a tube filled with 70% ethanol and labelled corresponding to the sample number of the host bat.

### Morphological identification of ectoparasites

Following field studies, ectoparasite samples collected from *E. dupreanum* and *R. madagascariensis* bats captured between February 2018 and November 2019 were examined under a standard light microscope (OMAX M8311) and subject to additional morphological assessment (hereafter, the ‘morphological data subset’). Under the microscope, different ectoparasite species were sorted morphologically and recounted into broad taxonomic categories (ticks, mites, fleas, and Nycteribiidae or Streblidae bat flies) following previously published taxonomic guides. These included general guides for bat ectoparasites broadly (65), specific references for bat mites (66,67) and specialized guides for Malagasy bat flies in both Nycteribiidae (51–53) and Streblidae (56,57) families. Where possible, ectoparasites were further categorized by genus and species, and bat flies were grouped by male and female sex. Photographs were taken of all observed genera of any ectoparasite taxa.

### Molecular identification of ectoparasites

Ectoparasite specimens from samples collected between February 2018 and November 2019 were exported to the University of Chicago for molecular identification. A random subset of bat flies corresponding to all three species observed during morphological study were selected for DNA extraction and barcoding (Table S2). For all selected specimens, DNA was extracted using the Zymo Quick-DNA 96 Plus Kit, following the manufacturer’s instructions and including the step for Proteinase K digestion. Following extraction, DNA quality was verified on a nanodrop, and high quality DNA samples were barcoded via amplification of the well-conserved cytochrome C oxidase subunit 1 gene (COI) (650bps), using LCO1490 and HCO2198 primers that have been previously published (68) and previously applied in molecular studies of Malagasy bat flies (49,54). All polymerase chain reactions (PCR) were conducted in 25μ*l* reaction mixtures containing 12.5μ*l* of GoTaq colorless master mix (Promega, Madison, WI), 8μ*l* of deionized water, 0.25μ*l* of each primer, and 3 μ*l* of extracted DNA. The amplification profile was 95^0^C for 2 min, followed by 35 cycles of 30s at 95^0^C, 30s at 49^0^C, and 30s at 72^0^C. A final extension step of 7 min at 72^0^C was realized. PCR products were separated by electrophoresis on 1% agarose gel, stained with SYBR Safe DNA gel stain (Invitrogen: S33102) and visualized under UV light. PCR products were purified and sequenced at the University of Chicago genomics core using both forward and reverse primers.

Following sequencing and phylogenetic analysis, we subsequently elected to additionally amplify the 18S gene of a subset of representative samples of each of the three bat fly species to compare against available reference sequences. Here, we used previously-published 1.2F and 7R PCR primers to target a 1600bp region of the 18S gene of extracted DNA samples (49,69,70) in a conventional (one-step) PCR protocol. All 18S PCR were conducted in 25μ*l* reaction mixtures following the same proportions as used for COI amplification, with the following amplification profile: 95^0^C for 4 min, followed by 40 cycles of 40s at 95^0^C, 30s at 57^0^C, and 60s at 72^0^C. A final extension step of 10 min at 72^0^C was realized. PCR products were separated by electrophoresis and purified, then sequenced at the University of Chicago genomics core following the same protocol used for COI barcoding.

Following barcoding, recovered sequences were manually curated in Geneious Prime 2022.1.1 (www.geneious.com): the 5’ and 3’ ends of all forward and reverse sequences were trimmed, low-support nucleotide calls were eliminated, and a consensus sequence was generated from paired forward and reverse sequences for all samples. Following cleaning, sequences were submitted to the Barcode of Life Database (BOLD) (71), then subsequently uploaded to NCBI GenBank.

### Climate data

Meteorological data used in these analyses were downloaded from NASA Earthdata Program using the Giovanni tool (https://giovanni.gsfc.nasa.gov/giovanni/) in raster format. We downloaded monthly temperature (^0^C), precipitation (mm), and diurnal humidity (% relative humidity, RH) data for all of Madagascar from January 2013 to December 2019 (Table S1). Monthly averages per year were calculated in R (v4.4.1) (72) for a 30-km buffer surrounding the Angavokely and Maromizaha roost sites (respectively for *E. dupreanum* and *R. madagascariensis*), for which we evaluated seasonal patterns in parasitism. The 30-km buffer was chosen as an appropriate spatial resolution to account for variation in the resolution of the different meteorological datasets, as well as to encompass the average distance a bat may travel in a typical foraging night.

### Data analysis

All analysis was conducted in R. All data and code can be accessed freely through our open-access Github repository: https://github.com/brooklabteam/Mada-Ectoparasites.

### Host-ectoparasite associations and correlations with field studies

Using the ‘bipartite’ package in R (73), we first constructed an alluvial plot to group host-ectoparasite relationships from the morphological data subset by bat species and associated ectoparasites categorized into Class, Order, Superorder, Family, and Genus.

Next, to evaluate the accuracy of our parasitological classifications in the field and evaluate whether field estimates of ectoparasite counts by taxonomic group across our entire 2013-2020 time series could be used to test ecological hypotheses, we compared morphological counts under the microscope of nycteribiid bat flies (*C. dubia* for *E. dupreanum* hosts and *E. madagascariensis* for *R. madagascariensis* hosts) against raw field counts of the same species. To this end, we used a simple linear regression to test the strength of association between nycteribiid count via microscopy in the laboratory and nycteribiid count in the field using the morphological data subset which reported both metrics. Because we only began reliably recognizing and recording counts of *M. wenzeli* several years into our time series, we did not attempt to compare field and laboratory counts of streblid parasites of *R. madagascariensis* but instead limited ecological analyses to nycteribiid bat flies only.

### Correlates of seasonal nycteribiid abundance using Generalized Additive Models (GAMs)

Because our comparison of field- and laboratory-derived nycteribiid counts suggested that we accurately estimated ectoparasite burden in the field (see ‘Results’), we carried out all subsequent analysis of seasonal patterns in parasitism using the field-derived dataset, which spanned from August 2013 – March 2020. Seasonal analyses reported in the main text were restricted to the subset of our data collected in central-east Madagascar (*E. dupreanum* roosts: Angavobe and Angavokely; *R. madagascariensis* roost: Maromizaha), where *E. dupreanum* were sampled in 11/12 months of the year (missing May only) and *R. madagascariensis* were sampled in all months of the year. We also report seasonal analyses for our northern Madagascar site (Ankarana National Park) in the Supplementary Materials, with the caveat that the temporality of these data are more limited: *E. dupreanum* were sampled in March-April and August-November in this site and *R. madagascariensis* in March and August-November. For both study regions, we further restricted seasonal analyses only to adult bats to allow for comparison of the impact of sex and body condition on nycteribiid bat fly abundance within each bat host species.

We used the ‘mgcv’ package in R (74) to construct generalized additive models (GAMs) aimed at identifying ecological correlates of the response variable of the abundance of nycteribiid ectoparasites infesting our two bat host species, separately for our two study regions. We first fit a series of Poisson GAMs to our data, evaluating the correlation of a suite of diverse predictor variables against the response variable of nycteribiid count, separately for *E. dupreanum* and *R. madagascariensis* bat hosts. We tested the hypothesis that ectoparasite abundance varied seasonally by allowing for a smoothing spline predictor of ‘day of year’ and a random effect of ‘sampling year’ in each GAM. We tested hypotheses that allowed for host sex-specific differences in the seasonality of parasitism (incorporating ‘bat sex’ in the smoothing spline ‘by’ term) vs. a composite seasonality across the two sexes. We also compared models which additionally incorporated random effect smoothing splines for the categorical variable of host bat sex and thinplate smoothing splines for mass: forearm residual (MFR), a measure of host body condition that we have previously shown to vary seasonally in these populations, tracking resource availability (61). We calculated MFR as the residual of the regression of log_10_ mass (in g) per log_10_ forearm length (in mm) for each separate sex (male vs. female) and species (*E. dupreanum* vs. *R. madagascariensis*) subset of our data. For all central-east analyses, we fixed the seasonal smoothing knots (‘k’) at 7 as recommended by the package author (74); for northern Madagascar data, we limited smoothing knots to 4 due to more limited seasonality in the data. In all cases, we modeled the ‘day of year’ smoothing spline as a cyclic cubic spline to force continuity from the end of one year to the beginning of the next. We compared all GAM formulations by Akaike Information Criteria (AIC) to determine the best fit to the data.

For the central-east study region, we additionally reran our GAM analysis on the morphological data subset, this time including the additional categorical predictor of bat fly sex, which we recorded during microscopy. As with the complete field datasets, we compared model fits by AIC and plotted significant predictor variables for both bat species and sex combinations. Too few individuals were morphologically evaluated from our northern Madagascar field site to allow for similar analysis for this region. Additionally, using the morphological data subset we carried out a two-sided student’s t-test comparing the mean abundance of male vs. female nycteribiids observed on *E. dupreanum* and *R. madagascariensis* bats for the two localities (central-east and north) surveyed.

### Cross-correlation analysis of nycteribiid association with climate

Because our GAMs indicated significant seasonality in nycteribiid abundance across our time series (see ‘Results’), we next evaluated the role of climate in driving this seasonal variation. To this end, we carried out cross correlation analysis in the R package ‘sour’ (75) to calculate the optimal lag between the mean nycteribiid bat fly (*C. dubia* for *E. dupreanum* and *E. madagascariensis* for *R. madagascariensis*) count per bat per month from 2013 - 2019 and the monthly average of our three climate variables (mean monthly diurnal humidity, mean monthly precipitation rate, and mean monthly temperature) for the corresponding locality (Angavokely cave for *E. dupreanum* and Maromizaha cave *R. madagascariensis*) across the same timespan. We considered monthly time lags up to one year by which climate variables preceded ectoparasite burden. Because our GAM analyses indicated significant seasonal deviations by host bat sex in ectoparasite burden (see ‘Results’), we calculated optimal lags to disparate ectoparasite burden time series for male and female host bats for the two species. Additionally, for visualization purposes, we summarized monthly averages across the entire study period for all three climate variables and for bat fly abundance for both bat hosts.

### Climate correlates of nycteribiid abundance using Generalized Linear Models (GLMs)

Following cross correlation analysis, we next constructed a composite dataset that included the three optimally lagged climate variables alongside the corresponding ectoparasite burden for each bat species and sex. Then, we compared a series of Poisson family generalized linear models (GLMs) to evaluate linear predictors of nycteribiid bat fly count, separately for *E. dupreanum* and *R. madagascariensis.* In addition to the three climate variables, models included predictor variables of bat sex and MFR. We compared model fits by AIC and reported the incidence rate ratio of all significant correlates in the top-performing model for each species.

### Phylogenetic analysis

Finally, using data generated from DNA barcoding, we constructed one COI and one 18S maximum likelihood (ML) phylogenetic tree comparing Madagascar bat fly (nycteribiid and streblid) DNA sequences with available reference sequences downloaded from NCBI and reported in previous studies (29,49,76,77). We rooted both phylogenies with *Drosophila melanogaster;* see Table S3 for NCBI accession numbers for all sequences (both new and reference) included in our phylogenetic analyses. For both COI and 18S phylogenies, we aligned sequences using the default parameters in MAFFT v7 (78,79), and checked alignments manually for quality control in Geneious Prime. We carried out all subsequent phylogenetic analyses on both a trimmed alignment of the conserved region of each gene (COI: 336bp; 18S: 391bp), in addition to an untrimmed version. As results were comparable across the two methodologies, we report results of only the untrimmed alignments here. All sequence subsets and alignment files (including trimmed versions) are available for public access in our GitHub repository: https://github.com/brooklabteam/Mada-Ectoparasites.

Following quality control, alignments were sent to Modeltest-NG (80) to assess the best fit nucleotide substitution model appropriate for our data. Both alignments (COI and 18S) were subsequently sent to RAxML-NG to construct the corresponding phylogenetic trees (81) using the best-fit nucleotide substitution model as estimated by Modeltest-NG (80). Following best practices outlined in the RAxML-NG manual, twenty ML inferences were made on each original alignment and bootstrap replicate trees were inferred using Felsenstein’s method (82), with the MRE-based bootstopping test applied after every 50 replicates (83). Bootstrapping was terminated once diagnostic statistics dropped below the threshold value and support values were drawn on the best-scoring tree. We plotted the resulting phylogenetic trees using the ggtree package in R (84).

## RESULTS

### Bat fly detection and host-parasite associations

From 2013-2020, we captured from 873 *E. dupreanum* bats (408 male, 465 female) and 862 *R. madagascariensis* bats (457 male, 405 female), which we surveyed for ectoparasites (nycterbiid and streblid bat flies, fleas, mites, ticks) (Table S1). Among those captured bats, we successfully counted, identified, and collected ectoparasites from 628 *E. dupreanum* (290 male, 338 female) and 831 *R. madagascariensis* (438 male, 393 female). We undertook detailed morphological analysis of a subset of ectoparasite samples collected from bats captured between February 2018 and November 2019 (*E. dupreanum:* 137 male, 214 female; *R. madagascariensis:* 241 male, 232 female). Among this morphological subset, we identified 264 (42%) *E. dupreanum* and 613 (74%) *R. madagascariensis* that hosted bat flies (family: Nycteribiidae or Streblidae); 114 (18%) *E. dupreanum* and 2 (<1%) *R. madagascariensis* that hosted fleas, 419 (68%) *E. dupreanum* and 660 (79%) *R. madagascariensis* that hosted mites, and 83 (13%) *E. dupreanum* and 36 (4%) *R. madagascariensis* that hosted ticks (**Fig. 1**). Simultaneous parasitism by multiple ectoparasite taxa was common on any individual bat: *E. dupreanum* were simultaneously parasitized by a mean 1.44 [95% confidence interval (CI): 1.36-1.51] different ectoparasite taxa (nycterbiids, streblids, fleas, mites, or ticks), while *R. madagascariensis* were parasitized by a mean 2.11 [95% CI: 2.05-2.17] ectoparasite taxa.

**Fig. 1.**
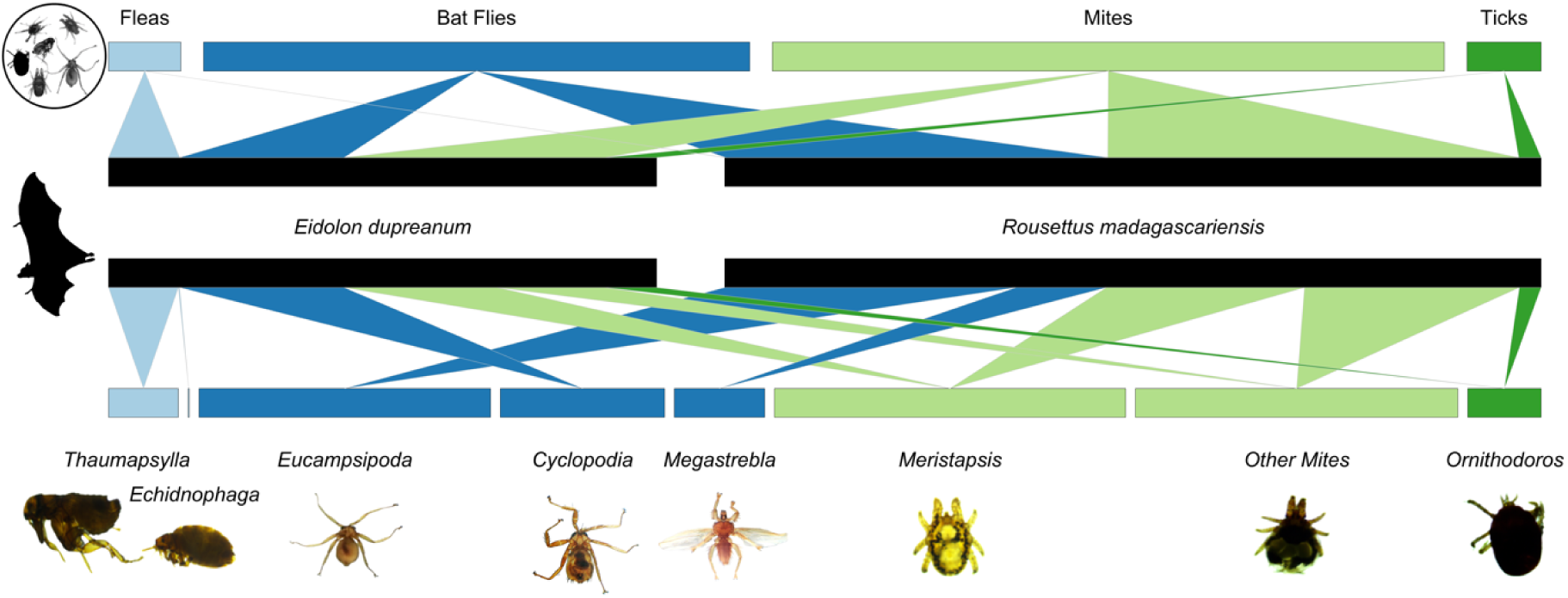
Alluvial plot showing bat host species (center) associations with broad ectoparasite clades (top) and genus-level classifications (bottom). Fleas and bat flies belonging to order Diptera are colored in shades of blue, while mites and ticks belong to class Arachnida (respectively in superorder Acariformes and Parasitiformes) are colored in shades of green. Images taken under the microscope at 40x magnification are shown below the names of the corresponding genera.

One species of nycteribiid bat fly, *C. dubia,* was identified on *E. dupreanum* bats. Both the nycteribiid *E. madagascariensis* and the streblid *M. wenzeli* were identified on *R. madagascariensis* bats. *E. madagascariensis* parasitism of *R. madagascariensis* was more frequent and occurred at higher abundance than *M. wenzeli* (Fig. 1). Most bat fleas infesting *E. dupreanum* were *Thaumapsylla* sp. previously reported on this host (4), while a few *Echidnophaga* sp. were also observed. Two fleas that keyed to family *Ischnopsyllidae* were also observed on two *R. madagascariensis* bats. Most mites parasitizing either bat host species belonged to the genus *Meristapsis,* though non-*Meristapsis* mites were also observed (66,67).

Ticks observed on both bat species keyed to the genus *Ornithorodos* in the soft-bodied tick family Argasidae (Fig. 1) (65). Downstream molecular assay is needed to confirm genus- and species-level identifications of flea, mite, and tick ectoparasites.

A linear regression comparing the laboratory recount of *C. dubia* on *E. dupreanum* and *E. madagascariensis* on *R. madagascariensis* against the raw field count observations demonstrated a highly significant positive correlation in both cases (**Fig. S1**; *C. dubia:* r = 0.92, p<0.001; *E. madagascariensis:* r=0.91; p<0.001), indicating that our field counts could be used representatively to explore broad seasonal patterns in our dataset.

### Correlates of seasonal nycteribiid abundance from GAMs

For both *E. dupreanum* and *R. madagascariensis* bat hosts in central-eastern Madagascar, the top-performing GAM to recover seasonal nycteribiid abundance included a cyclic smoothing spline predictor by ‘day of year’ with a ‘by’ term of ‘bat sex’ allowing for disparate seasonal trends of parasitism for male vs. female bat hosts (**Fig. 2**; Table S4). For both host species, the top-performing model included a predictor of MFR, though this variable only demonstrated partial significance in *R. madagascariensis* models (Fig. 2B,D). The abundance of *C. dubia* on *E. dupreanum* peaked in ∼late May/early June for female bats (preceding the onset of the gestation period) and late June for male bats (at the onset of the nutritionally scarce dry season). The abundance of *E. madagascariensis* on *R. madagascariensis* peaked in late February/early March for females (∼5 months preceding gestation) and March for males during the resource-abundant wet season. Despite improving overall model fit, MFR showed no significant variation with bat fly abundance for *E. dupreanum* (Fig. 2B). For *R madagascariensis*, extremely low MFR values were associated with lower bat fly burden, and high MFR values were associated with slightly elevated bat fly burden (Fig. 2D).

**Fig. 2.**
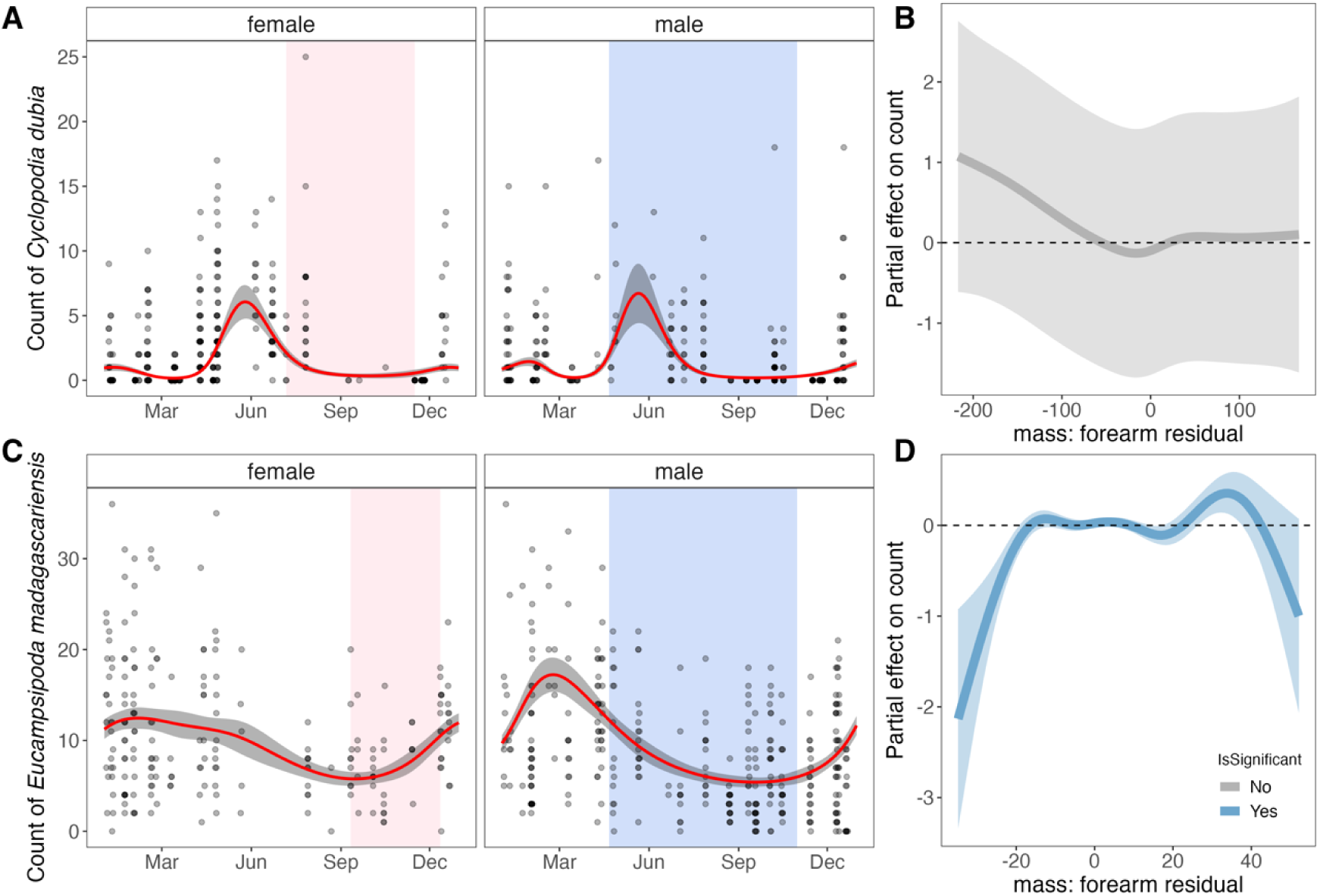
Seasonal variation in the abundance of Nycteribiidae bat flies counted on (**A, B**) *E. dupreanum* and (**C,D**) *R. madagascariensis* bats captured at roost sites in central-eastern Madagascar (respectively, Angavobe/Angavokely and Maromizaha caves). Panels (A) and (C) show seasonal ectoparasite count predictions (red line) from best-fit GAMs for male and female bat hosts of each species, with 95% CI by standard error shaded in gray. Translucent background points in black correspond to raw data across all years of the study (2013-2019). Pink background shading corresponds to the gestation period for each species from (61), while blue background shading corresponds to the nutritionally deficient dry season for the region. Panels (B) and (D) show partial effect (y-axis) of bat host mass: forearm residual, respectively for *E. dupreanum* and *R. madagascariensis,* on bat fly count. Solid lines (gray for non-significant effects; blue for significant effects) correspond to mean effects, with 95% CIs by standard error shown in translucent shading.

GAMs demonstrated similar results when refit to the Feb 2018-Nov 2019 morphological data subset (**Fig. S2;** Table S4). Inclusion of additional categorical predictors of bat host sex and bat fly sex improved model performance against the morphological data subset, but partial effects for these predictors were not significant. For GAMs fit to the morphological data subset, MFR had a significant effect on bat fly count for both *E. dupreanum* and *R. madagascariensis,* largely recapitulating trends from the full field dataset (Fig. 2B,D). For *E. dupreanum* hosts, low MFR was associated with high bat fly burden and high MFR with lower bat fly burden (Fig. S2B); patterns were reversed for *R. madagascariensis* hosts, where low MFR was again associated with low bat fly burden, and high MFR was associated with higher bat fly burden (Fig. S2D).

A student’s t-test demonstrated no significant difference in the average count of male vs. female *C. dubia* recovered on *E. dupreanum* hosts (p=0.015) or male vs. female *E. madagascariensis* recovered on *R. madagascariensis* hosts (p=0.07) in the central-eastern data, where sampling was representative across the entire calendar year (**Fig. S3**). We did observe significantly higher mean count of male vs. female nycteribiids for both *E. dupreanum* and *R. madagascariensis* hosts in the morphological data subset from northern Madagascar (p<0.001 in both cases), where sampling was restricted to just the dry season months of the year (Fig. S3).

For the northern Madagascar field-derived dataset, the best-fit model for both *E. dupreanum* and *R. madagascariensis* hosts included predictor variables of host sex-specific seasonal smoothing splines, in addition to MFR and a random effect of host bat sex (**Fig. S4**; Table S4). Though the seasonal duration of these data were more limited, we estimated a slightly later peak in nycteribiid burden as compared with central-eastern study sites, with highest abundance observed in ∼late August/early September for female *E. dupreanum* and September for males and in ∼late September/early October for female *R. madagascariensis* and October for males (Fig. S4). The limited sampling window of our data does not preclude the possibility of a second peak in bat fly abundance early in the calendar year. Models fit to data from northern Madagascar recapitulated patterns observed in central-east Madagascar for ectoparasite relation to MFR: no significant correlations were observed for *E. dupreanum,* though patterns trended to higher bat fly load in low MFR individuals (Fig. S4B). Significant trends were observed for *R. madagascariensis,* again showing the opposite pattern, with lower bat fly burden in bats with the lowest MFR (Fig. S4E). Though inclusion of bat sex improved model fits to this northern Madagascar data subset, no significant partial effects by sex were observed (Fig. S4C,E).

### Cross-correlation analysis of nycteribiid association with climate

Because GAM analyses indicated significant host sex-specific seasonality in bat fly burden for both bat species, we next investigated the correlation between site-specific climate variables and seasonal variation in nycteribiid bat fly count for *E. dupreanum* and *R. madagascariensis.* We first plotted the monthly average of three key climate variables (daily humidity, precipitation rate, and temperature) for each roost site (Angavokely and Maromizaha caves), as compared to the monthly average bat fly count per bat for the two species (**Fig. 3**). We observed a substantial lag between monthly peaks in precipitation and temperature climate variables and the corresponding peak in ectoparasite abundance for both localities studied. We next quantified these lags using cross correlation analysis of the monthly average for each climate variable per year from 2013-2019, compared against the time series of bat fly abundance on male and female bats of both species (**Fig. S5**; Table S5A). For *E. dupreanum,* the cross correlation between climate variable and bat fly abundance was maximized at no lag for humidity and abundance on male bats and a 5 month lag for females; at a 4 month lag for precipitation and abundance on male bats and a 5 month lag for females; and at a 3 month lag for temperature and abundance on male bats and a 6 month lag for females (Fig. S5; Table S5). For *R. madagascariensis,* the cross correlation between climate and bat fly time series for both male and female bats was maximized at no lag for humidity and one month for both precipitation and temperature time series (Fig. S5B).

**Fig. 3.**
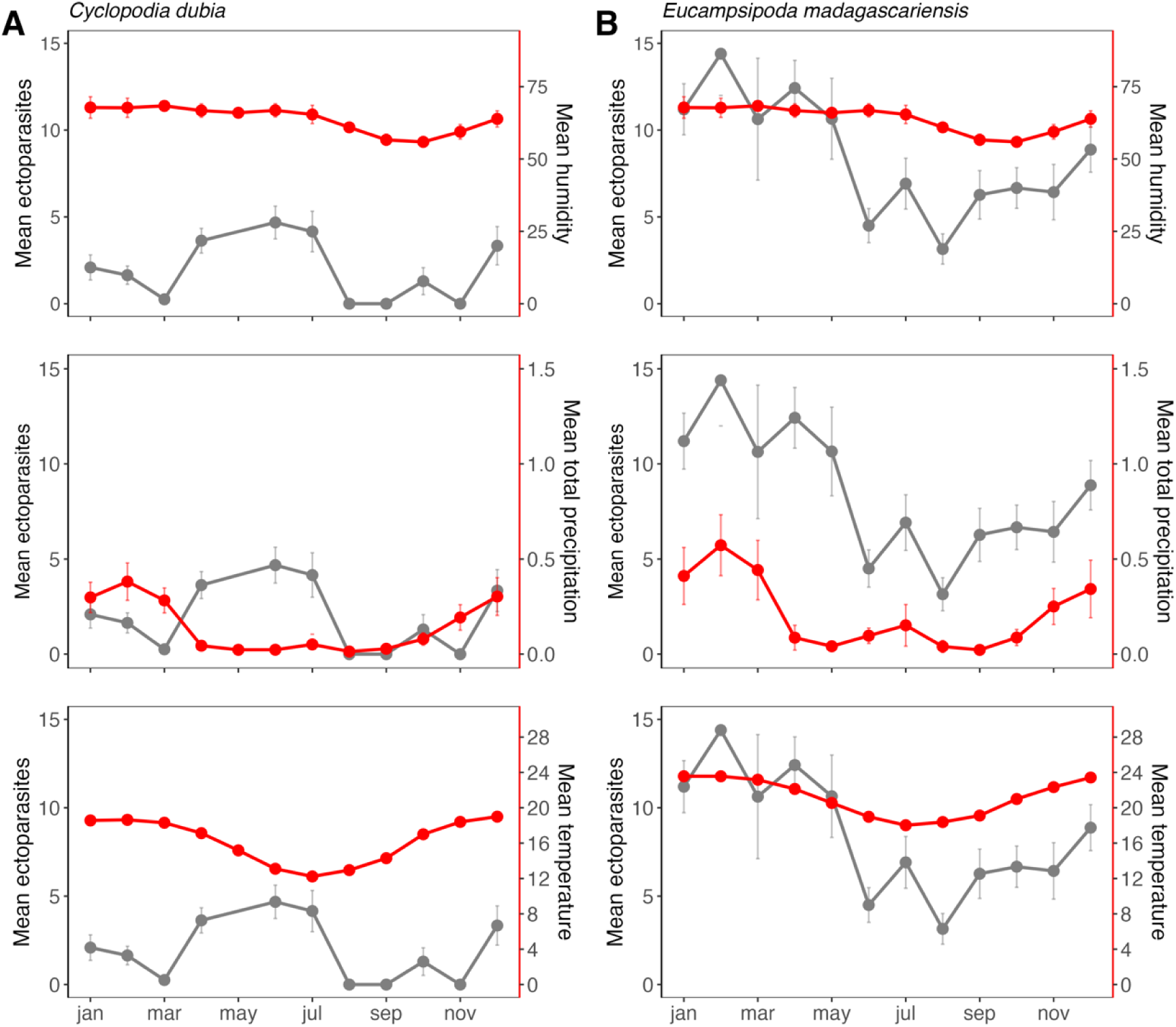
Mean monthly nycteribiid count per bat for (**A**) *C. dubia* parasitism of *E. dupreanum,* across the full 2013-2020 time series for Angavokely roost site (gray lines and points; left y-axis), as compared with climate variables of monthly averages of (horizontal panels) diurnal humidity (% relative humidity), total precipitation (mm), and temperature (^0^C) for the same region (red lines and points; right y-axis). (**B**) *E. madagascariensis* parasitism of *R. madagascariensis,* mirroring the same structure in (A), for the Maromizaha roost site. 95% CIs by standard error are shown for both ectoparasite and climate data.

### Climate correlates of nycteribiid abundance from GLMs

Using the optimally lagged climate time series, we next identified linear predictors of nycteribiid bat fly burden across our field-derived dataset for both bat hosts (**Fig. 4**). For *C. dubia* abundance on *E. dupreanum,* the best-fit GLM included all predictor variables tested: all three climate variables (optimally lagged by bat host sex), in addition to bat sex and MFR (Fig. 4A). Lagged precipitation and temperature were the most influential variables contributing to overall model performance (Fig. 4A). All three climate variables were positively correlated with bat fly burden, while male bat sex was negatively correlated with bat fly abundance, and the effect of MFR was not significant (Fig. 4B). For *E. madagascariensis* abundance on *R. madagascariensis,* the best-fit GLM included all predictor variables tested except for MFR (Fig. 4C). Here, lagged temperature was the most influential variable contributing to overall model performance (Fig. 4C). As with the *E. dupreanum* model, all climate variables in the *R. madagascariensis* model were positively correlated with bat fly burden, and male bat sex was negatively associated with bat fly count (Fig. 4D).

**Fig. 4.**
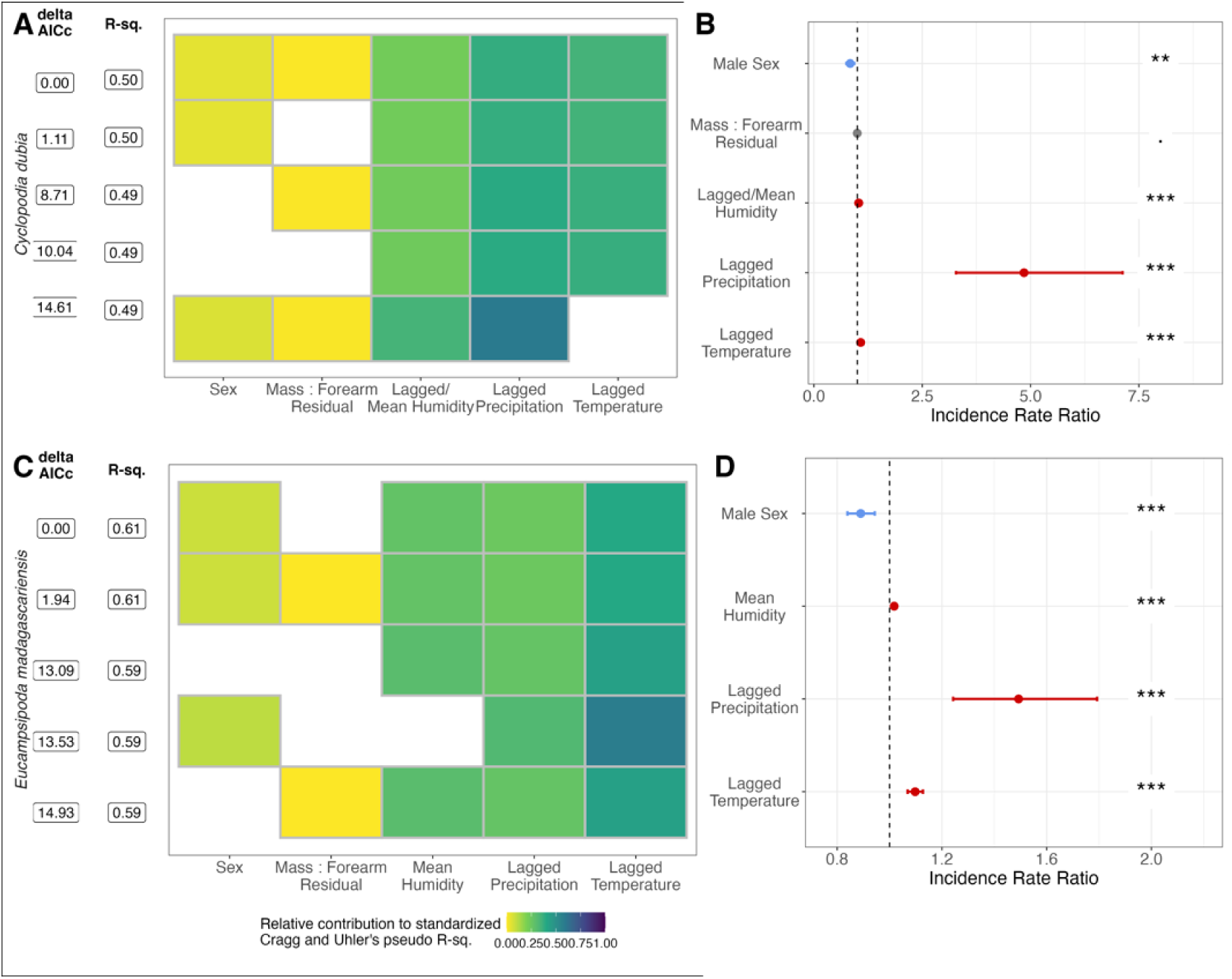
Influence of climate and demographic variables on the abundance of nyteribiid bat fly count for (**A,B**) *C. dubia* abundance on *E. dupreanum* and (**C,D**) *E. madagascariensis* abundance on *R. madagascariensis.* (**A,C**) Top five GLMs using optimally-lagged climate variables to predict bat fly abundance, ranked by δAICc, for *E. dupreanum* and *R. madagascariensis* bat hosts. Rows represent individual models and columns represent predictor variables. (**B,D**) Incidence rate ratios of each linear predictor from top-fit models shown respectively in (A,C). Significant positive correlates are colored red, significant negative correlates are colored blue, and insignificant correlates are colored grey. 95% CIs by standard error are shown as horizontal error bars.

### Phylogenetic inference

Nycteribiid and streblid sequences from bat flies of Malagasy *E. dupreanum* and *R. madagascariensis* recovered from COI and 18S DNA barcoding were deposited to GenBank under accession numbers listed in Table S2 (43 COI and 12 18S sequences), then aligned with available reference sequences for phylogenetic analysis (Table S3). Modeltest-NG (80) identified the best-fit nucleotide substitution model as GTR+I+G4 for the COI phylogeny and TIM2+I+G4 for the 18S phylogeny. Correspondingly, RAxML-NG (81) recovered similar topologies for both ML phylogenies (**Fig. 5**; **Fig. S6-S7**). COI sequences recovered from *C. dubia* parasitizing *E. dupreanum* clustered with previously-published sequences from this same species in a monophyletic clade with other *Cyclopodia* spp. identified from other Pteropodidae fruit bat hosts (Fig. 5; Fig. S6); our 18S *C. dubia* sequences represent the first *Cyclopodia* spp. contributions for this gene to GenBank (Fig. S7). Likewise, both COI and 18S sequences recovered from *E. madagascariensis* parasitizing *R. madagascariensis* clustered with previously-published sequences from this species. The *Eucampsipoda* spp., including *E. madagascariensis,* formed a different monophyletic clade within the Nycteribiidae family, with each disparate parasite species resolving into disparate subclades associated with a unique pteropodid fruit bat host species (Fig. 5; Fig. S6-S7).

**Fig. 5.**
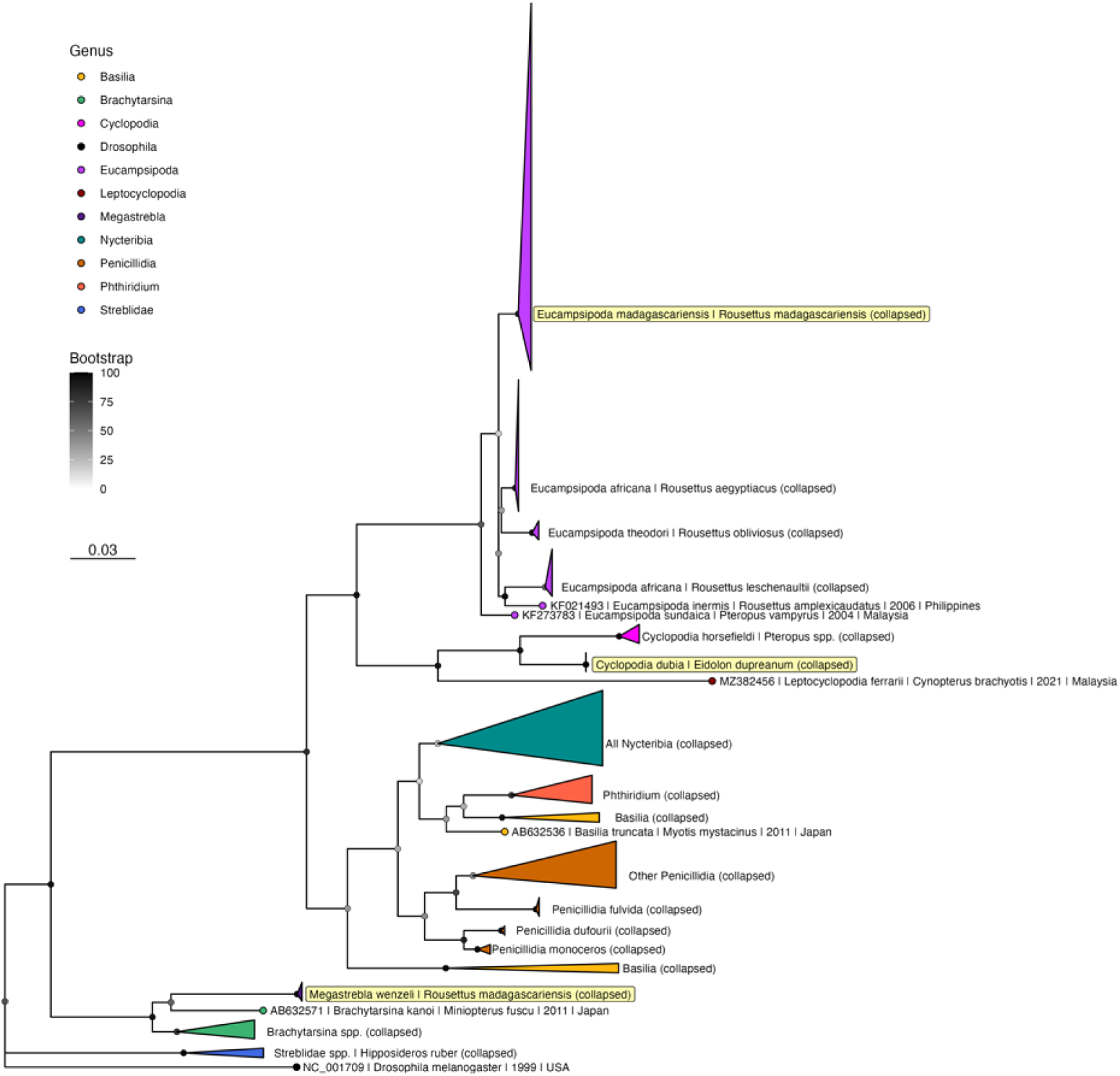
Maximum likelihood phylogeny of COI ectoparasite sequences from untrimmed alignment (RAxML-NG, GTR+I+G4) (81). Bootstrap support values computed using Felsenstein’s method (82) are indicated by shaded circles on each node, corresponding to legend. Sequences are collapsed into single species clades, or where indicated, clades by genera for ease of visualization; see Fig. S6 for full phylogeny with individual sequences labeled. Tip shapes are colored by genera, and tip labels for the three Madagascar clades (*E. madagascariensis, C. dubia, M. wenzeli*) are highlighted in yellow. Tree is rooted in *Drosophila melanogaster,* accession number NC_001709. Branch lengths are scaled by nucleotide substitutions per site, corresponding to the scalebar shown.

Both COI and 18S sequences recovered from *M. wenzeli* parasitizing *R. madagascariensis* represent the first molecular record of this streblid ectoparasite available on GenBank (Fig. 5; Fig. S6-S7). While to our knowledge no COI reference sequences are currently available for the *Megastrebla* genus, our 18S sequences from *M. wenzeli* clustered within a monophyletic clade of previously-reported *Megastrebla* spp. sequences recovered from parasites of other pteropodid fruit bats (Fig. S7) (29).

## DISCUSSION

We report diversity and seasonality in ectoparasite infestation of two species of endemic Malagasy fruit bat, *E. dupreanum* and *R. madagascariensis.* Both bat species were observed to be frequently co-parasitized by a suite of diverse ectoparasite taxa, most commonly bat flies in the family Nycteribiidae: *C. dubia* for *E. dupreanum* and *E. madagascariensis* for *R. madagascariensis,* consistent with previously published work (49,51–54). In addition, we report the first-ever molecular records documenting parasitism of *E. dupreanum* bats by the streblid ectoparasite, *M. wenzeli;* sequences recovered from our study place this parasite in a monophyletic clade of Old World streblids including previously-reported sequences for *M. nigriceps* and *M. parvior,* streblid bat flies collected from *Eonycteris spelaea* fruit bats in Malaysia (29). Our morphological observations of *M. wenzeli* are consistent with prior records describing this ectoparasite in Madagascar (56,57,59). In addition to bat flies, we reconfirmed previous reports of *E. dupreanum* parasitism by *Thaumapsylla* sp. fleas (4), in addition to parasitism of both *E. dupreanum* and *R. madagascariensis* parasitism by mites and ticks (4,49). Additional molecular studies will be needed to confirm species-level identity of fruit bat ectoparasites beyond bat flies in the superfamily Hippoboscoidea.

The bulk of our analyses centered on understanding seasonal variation in nycteribiid parasitism of the two fruit bat hosts. Previous work corresponding to this theme has been published for *E. madagascariensis* parasitism of *R. madagascariensis* in northern Madagascar (Ankarana National Park) (59,60). To our knowledge, our study is the first to document seasonal patterns of parasitism for *C. dubia* on *E. dupreanum,* as well as the first to document these patterns for either nycteribiid in central-eastern Madagascar (Districts of Manjakandriana and Moramanga), which has a cooler climate profile than the north. In general, our seasonal analyses in northern Madagascar mirrored those previously reported for *E. madagascariensis* parasitism of *R. madagascariensis:* we observed highest abundance of ectoparasite load per bat during the regional dry season (∼September), though our observations were too limited during the wet season (December – April) to rule out the possibility of a second annual peak. Some recent evidence suggests that *R. madagascariensis* may undergo two annual breeding seasons in northern Madagascar (85), a pattern previously reported for sister species *R. aegyptiacus* in more tropical localities on the African continent (86,87). As hormonal changes associated with reproduction are known to impact the seasonality of ectoparasite burden in other host-parasite systems (88–90), including bat systems (91–93), these reproductive changes may influence ectoparasite seasonality in our Madagascar system, as well. *C. dubia* parasitism of *E. dupreanum* in our northern Madagascar locality also peaked in September, though limited data in wet season months again precluded inference earlier in the year.

At our well-sampled central-eastern Moramanga site, we observed only one peak in nycteribiid burden for *R. madagascariensis,* towards the end of the wet season (March) for this locality; in related studies, we have only observed a single annual gestation period between September and December for *R. madagascariensis* in the same site (61,94). Also in our central-eastern study region, we observed a single peak in ectoparasite burden for *C. dubia* parasitism of *E. dupreanum,* here preceding the onset of the female gestation period for this bat species, at the start of the dry season (June) in this region. We note that, while we group both the Angavokely roost for *E. dupreanum* and Maromizaha roost for *R. madagascariensis* within the central-eastern region of Madagascar, these sites are located over 60km apart (Table S1), and mean monthly temperatures were on average just under 5^0^C cooler in Angavokely vs. Maromizaha across our study period (Fig. 3). This suggests different climatic influences on both the reproductive calendar for the bat hosts and the seasonality of ectoparasite burden in the two localities. Our GLM analyses highlight an important role for climate, particularly precipitation and temperature, in driving seasonality in ectoparasite burden, whether directly through impacts on ectoparasite physiology or indirectly through modulation of bat host physiology or both. Clearly, seasonal patterns in ectoparasite burden are more comparable between the two bat species when sampled in the exact same northern Madagascar locality (Ankarana National Park) than when sampled in climatically different sites in central-eastern Madagascar. Nonetheless, despite the clear influence of climate, one key finding from our study is the repeated support we recovered across both study sites and both bat host species for sex-specific differences in the seasonality of ectoparasite burden, with ectoparasite burden on female bats always preceding that for males of the same species in the same locality. These patterns suggest that, independent of climate, seasonal differences in bat physiology, likely related to reproduction, are important drivers of ectoparasite burden.

In addition to seasonality, our analyses of the impact of bat host body condition (MFR) on ectoparasite load mirrored previous reports for *E. madagascariensis* on *R. madagascariensis* (60): we found higher parasite loads in individuals with better body condition (higher MFR), which, as previously hypothesized, could be related to larger surface area available for nycteribiid infestation in these relatively small (∼60g) fruit bats. We consistently observed the inverse trend, with low host MFR association with higher nycteribiid burden for larger (∼250g) *E. dupreanum* bats, perhaps due to immunocompromising effects of host nutrition. It is possible that *R. madagascariensis* fall below a certain size threshold below which available surface area scales positively with ectoparasite burden, while above this threshold, the effects of host physiology and immunology prevail. Further research at the minimum and maximum size distributions for these different species will be needed to parse these divergent trends.

In contrast to a previous study in northern Madagascar which identified higher nycteribiid parasitism intensity on adult male vs. female *R. madagascariensis* (59), we found that, after controlling for climate predictors, nycteribiid abundance was lower on male vs. female *R. madagascariensis* and *E. dupreanum* bats in our central-eastern Madagascar sites (Fig. 4). No significant effects of host bat sex were observed for either species in the more limited northern Madagascar dataset. Our findings in central-eastern Madagascar are consistent with previous reports of ectoparasite preference for female bat hosts in other systems (95,96) and suggest that previous reports in northern Madagascar may reflect seasonal biases in data collection, as hypothesized by the study authors. This prior study in northern Madagascar additionally identified a significant male sex bias in the *E. madagascariensis* ectoparasites themselves (59). We observed a similar bias in our northern Madagascar site, for which morphological observations were only conducted during the dry season (Fig. S3). As our more complete seasonal time series in central-eastern Madagascar showed no significant differences in sex distribution for either *C. dubia* or *E. madagascariensis* bat flies, we hypothesize that this previous report likely also reflects seasonal bias in the sampling. Indeed, previous reports in the literature suggest that seasonal variation in the sex ratio of nycteribiid bat fly populations may be common (97–99). Further field study is needed to more clearly delineate these patterns for *C. dubia* and *E. madagascariensis*.

Our study has several limitations, most obviously our reliance on publicly-available coarse-scale climate data in lieu of direct climate records collected from the interior of bat cave roosts. Previous studies have demonstrated critical impacts of microclimate differences in bat roosts on seasonal dynamics in ectoparasite communities (100,101). As we observed clear differences in the seasonality of ectoparasite burden across study sites within the same broad geographic region, future work will greatly benefit from more careful study of local climate. In addition, our more limited seasonal sampling of northern Madagascar localities (due to access challenges in the peak rainy season) precludes some comparisons between our two study regions; equally intensive seasonal study of northern Madagascar sites will offer additional insight in the future. We focused the bulk of our ecological analyses on seasonal variation in the abundance of nycteribiid bat flies; future work should attempt to carry out similar investigations into the ecology of *M. wenzeli* parasitism, in addition to the several other ectoparasite taxa observed during our field and laboratory studies. Finally, confirmation of species-level identity of non-bat fly ectoparasites of *E. dupreanum* and *R. madagascariensis* using molecular techniques is a major research priority.

## CONCLUSIONS

Ectoparasites of bats, including nycteribiid and streblid bat flies, fleas, mites, and ticks, can play important roles in the transmission of microparasitic infections. Here, we describe the diversity of ectoparasite burden for two species of Malagasy fruit bat, *E. dupreanum* and *R. madagascariensis,* expanding the existing molecular record to include streblid bat fly ectoparasites of *R. madagascariensis.* We additionally highlight seasonal variation in nycterbiid burden for these two bat hosts, which mirrors seasonal variation in nutritional resource availability and the reproductive calendar across northern and central-eastern Madagascar. As bats are important reservoirs for several highly virulent zoonotic microparasites (1), understanding ecological patterns of bat parasitism is of critical public health importance. As ectoparasites can cause negative fitness impacts on their hosts (102,103), our work is additionally informative for conservation efforts for these two Pteropodidae fruit bats, both ranked as ‘Vulnerable’ on the IUCN Red List of Threatened Species (104).

## DECLARATIONS

### Ethics approval and consent to participate

Not applicable.

### Consent for publication

Not applicable.

### Availability of data and materials

The datasets generated and/or analysed during the current study are attached as supplementary tables to this article and also available in our open-access GitHub repository: https://github.com/brooklabteam/Mada-Ectoparasites.

## Authors’ contributions

AFA, SA, HCR, SG, and CEB collected the field data in part with a large project overseen by CEB, JMH, PD, and VL. AFA and SA carried out microscopy on field-collected ectoparasite samples, with support from HCR and SG. AFA, SA, and GK conducted DNA barcoding of field-collected ectoparasite samples, with support from KIY and CEB. AFA analyzed the resulting data in R with support from TML, KIY, AA, and CEB. AFA and CEB wrote the first draft of the manuscript. All authors read and approved the final manuscript.

## Acknowledgements

We thank Anecia Gentles, Kimberly Rivera, and Fifi Ravelomanantsoa for help in the field and lab. We acknowledge the Virology Unit at the Institut Pasteur de Madagascar for logistical support, and we thank the Mention of Zoology and Animal Biodiversity at the University of Antananarivo and the Madagascar Ministry of the Environment and Sustainable Development for providing research and export permits. We thank the Brook lab at the University of Chicago for helpful contributions to the manuscript. This work was completed in part with resources provided by the University of Chicago’s Research Computing Center.

