## Supplementary figures and images for "Diversity and seasonality of ectoparasite burden on two species of Madagascar fruit bat, *Eidolon dupreanum* and *Rousettus madagascariensis*"

### Fig. S1

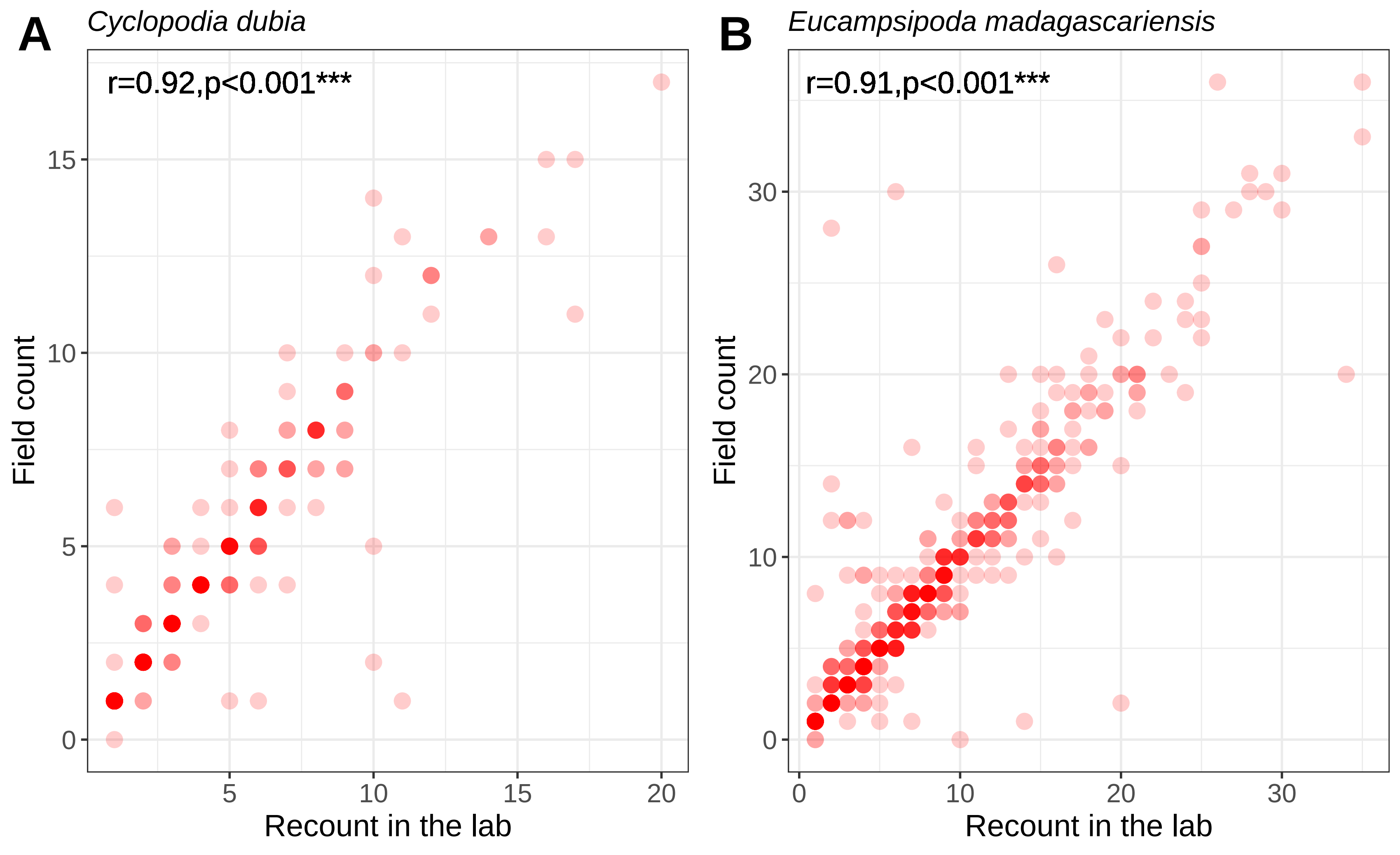

### Fig. S2

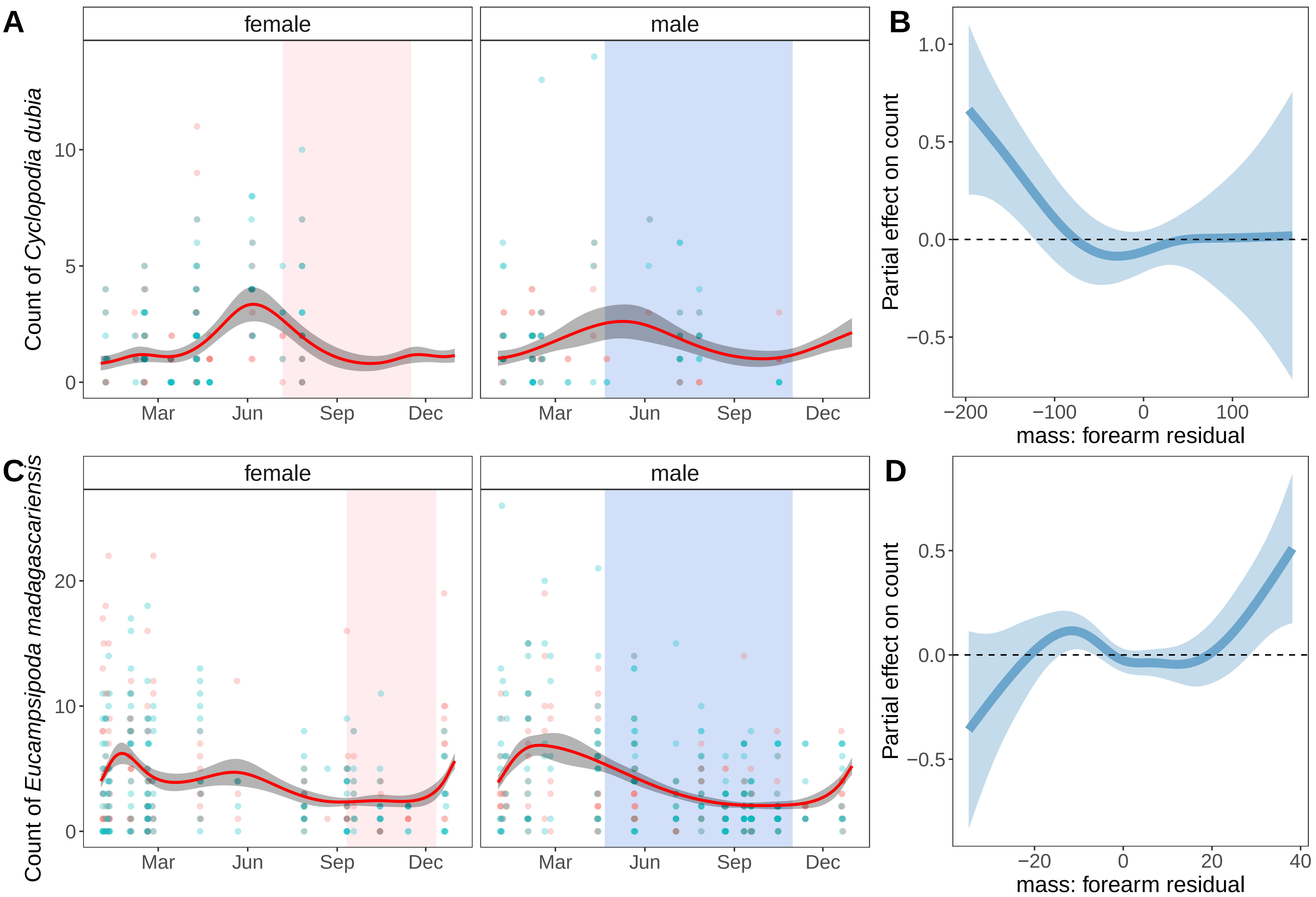

### Fig. S3

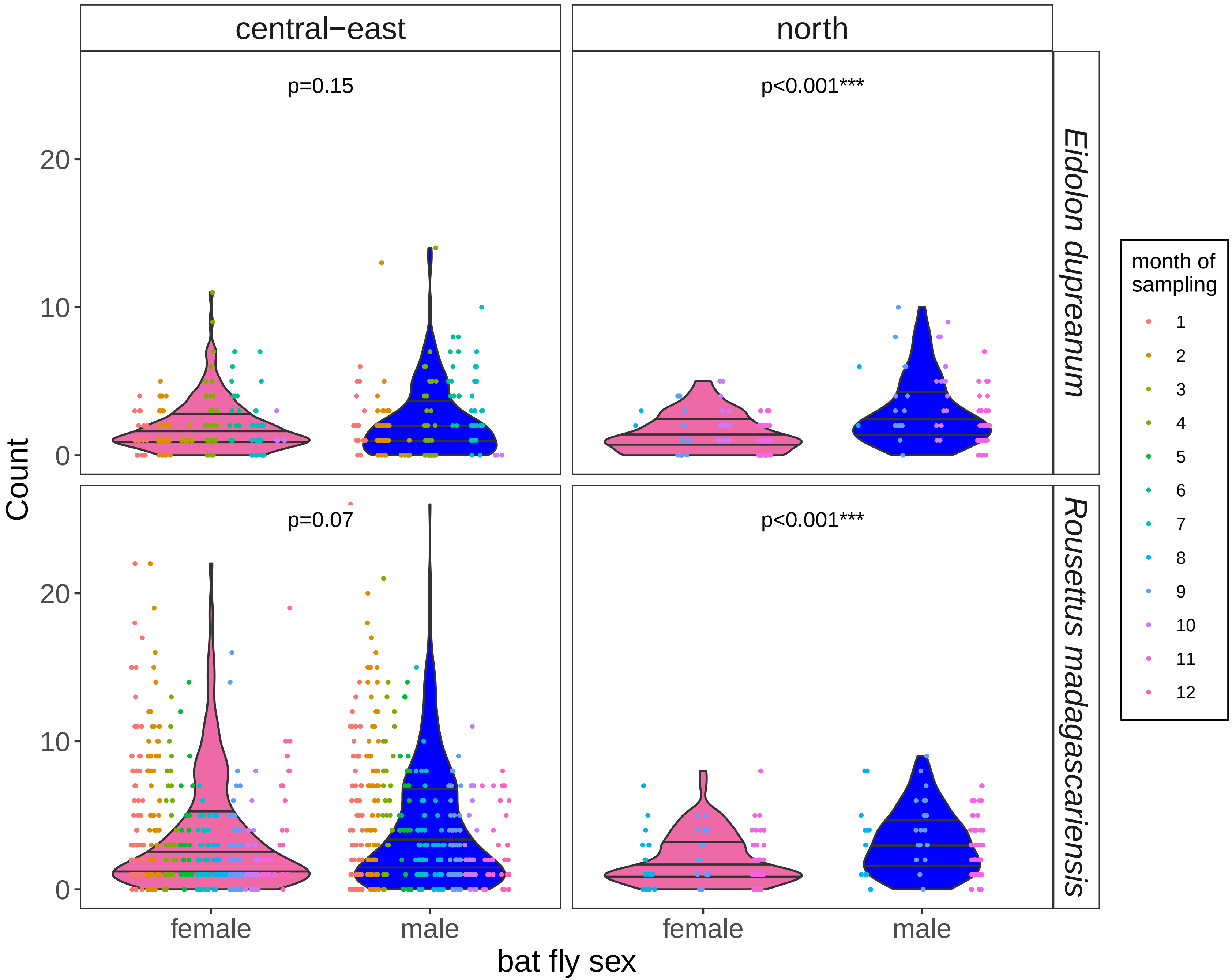

### Fig. S4

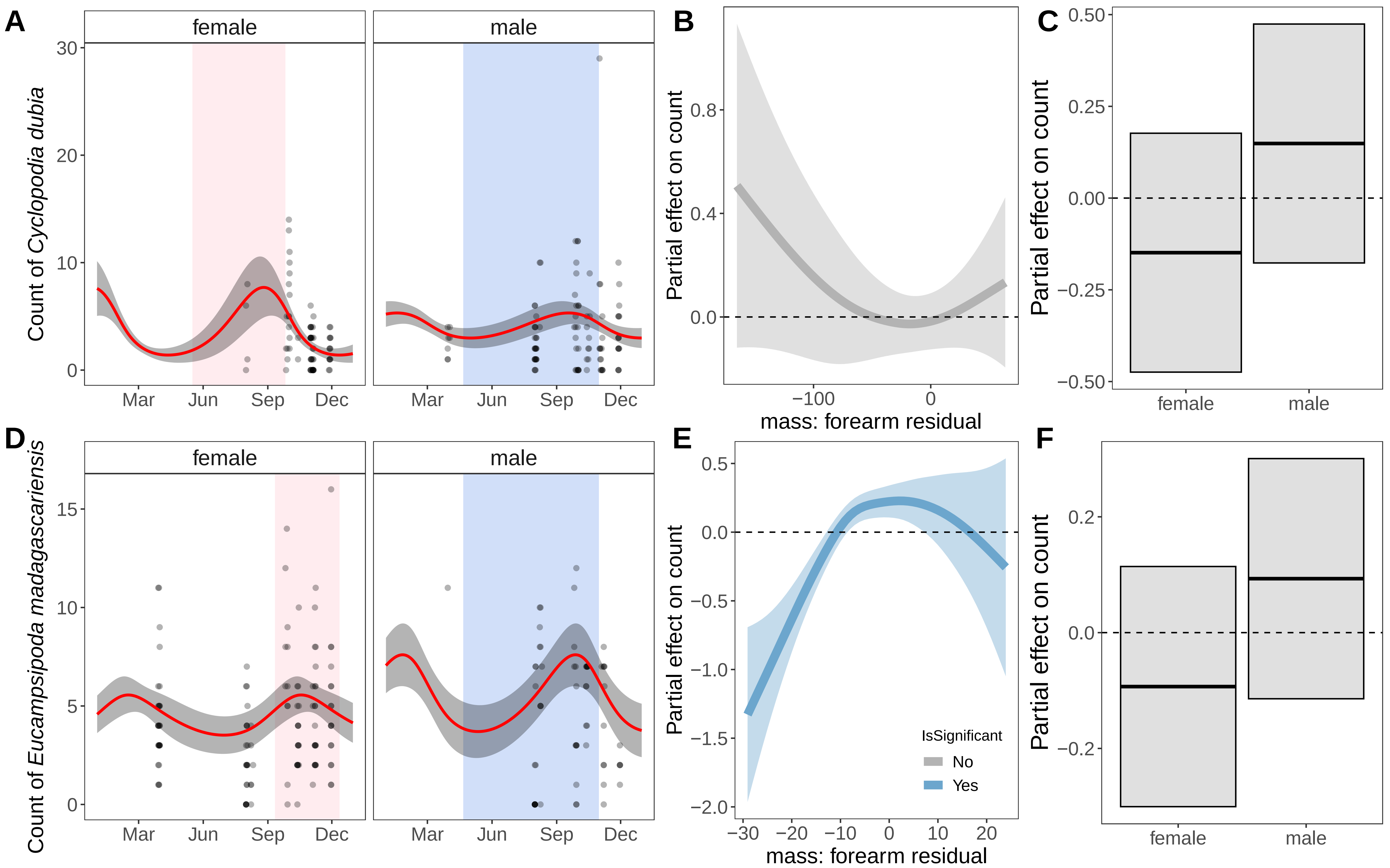

### Fig. S5

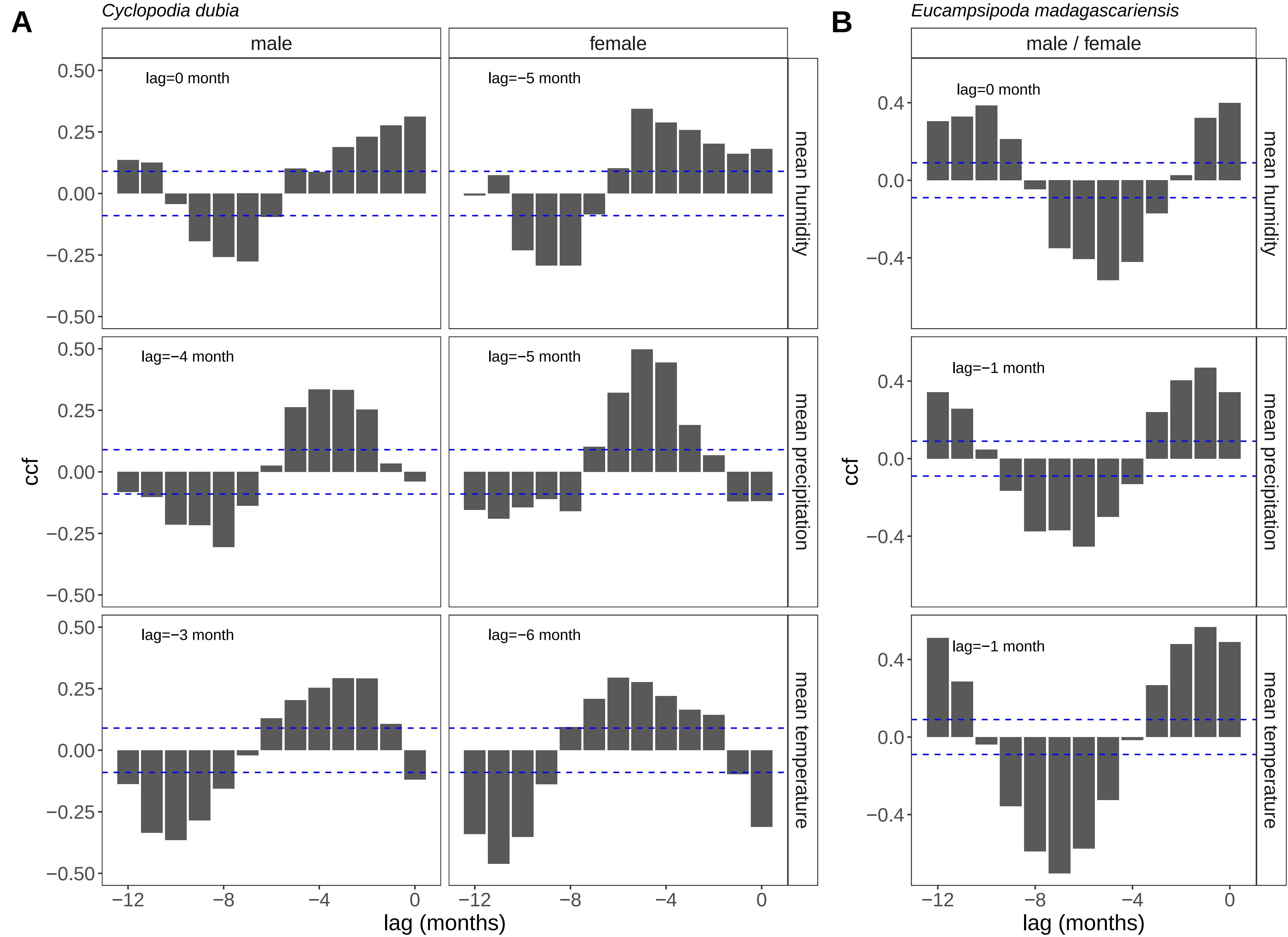

### Fig. S6

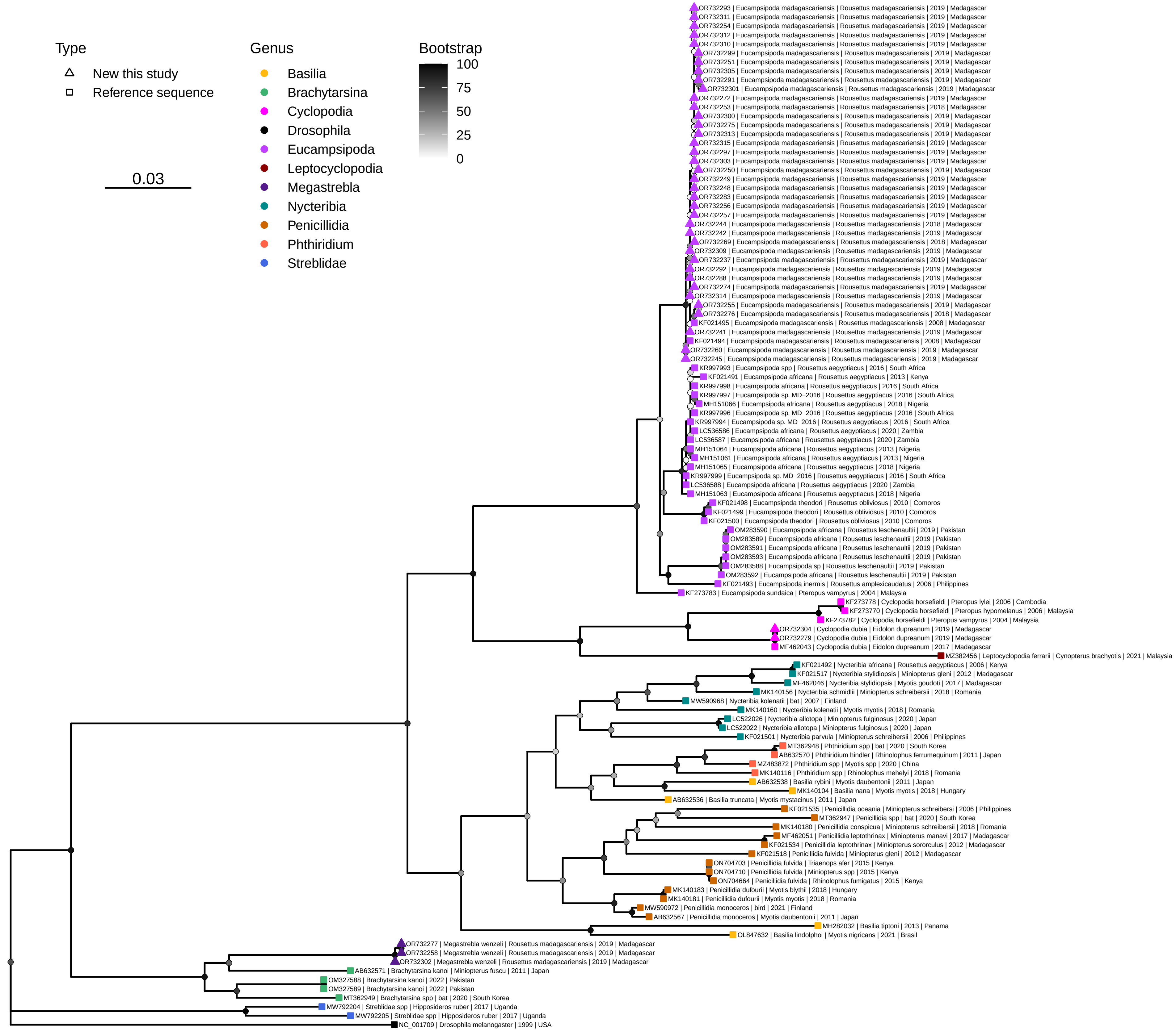

### Fig. S7

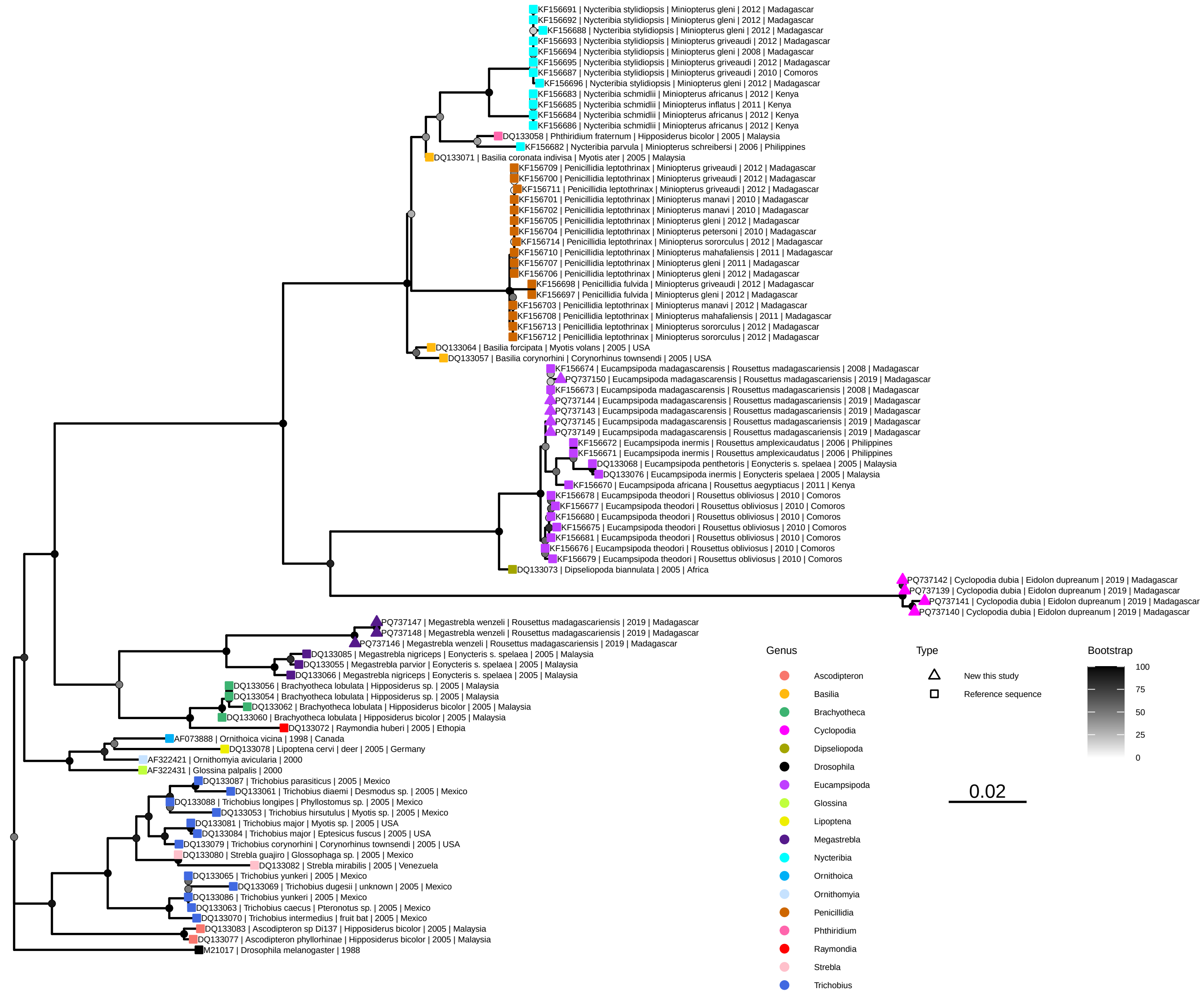
