## Supplementary material for "Diversity and seasonality of ectoparasite burden on two species of Madagascar fruit bat, *Eidolon dupreanum* and *Rousettus madagascariensis*": Fig. S

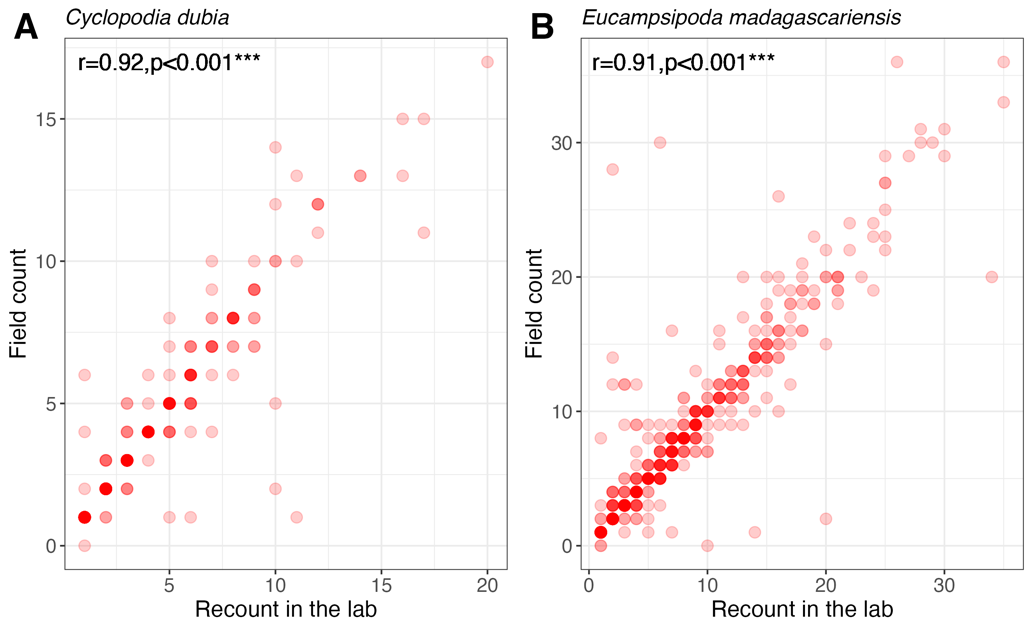

**Fig. S1.** Correlation analysis of morphological count in the laboratory (x-axis) vs. raw field counts of nycteribiid abundance on captured bats between Feb 2018 and Nov 2019, for (**A**) *C. dubia* parasitism of *E. dupreanum* bat hosts and (**B**) *E. madagascariensis* parasitism of *R. madagascariensis* bat hosts. Data are shown as translucent red points. The correlation coefficient (*r*) and corresponding p-value for each regression are listed in the top-left of each panel.

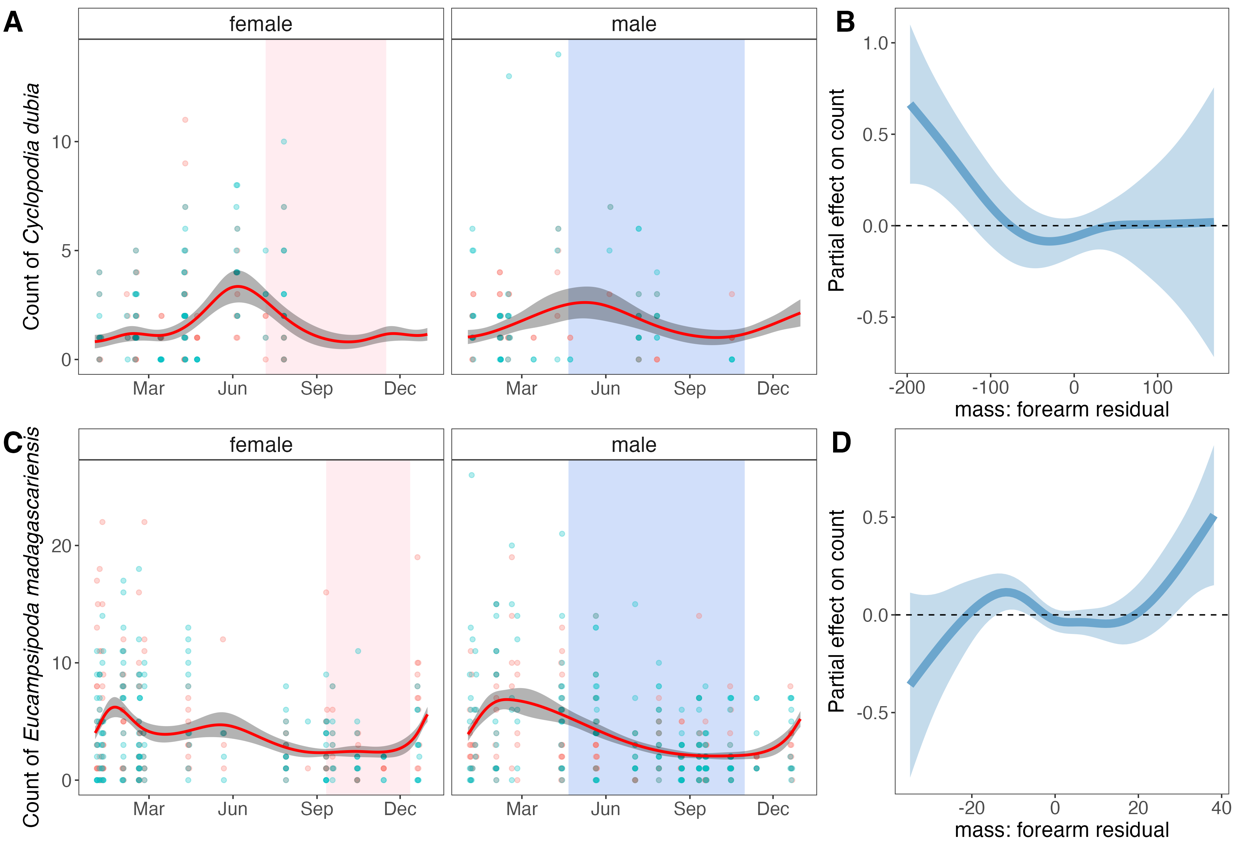

**Fig. S2.** Seasonal variation in the abundance of nycteribiidae bat flies counted on (**A, B**) *E. dupreanum* and (**C,D**) *R. madagascariensis* bats captured at roost sites in central-eastern Madagascar (respectively, Angavobe/Angavokely and Maromizaha caves) from morphological data subset. Panels (A) and (C) show seasonal ectoparasite count predictions (red line) from best-fit GAMs for male and female bat hosts of each species, with 95% CI by standard error shaded in gray. Translucent background points correspond to raw data across all years of the data subset (Feb 2018 – Nov 2019), with male bat flies represented in blue and female bat flies in pink. Pink background shading corresponds to the gestation period for each host bat species from (1), while blue background shading corresponds to the nutritionally deficient dry season for the region. Panels (B) and (C) show partial effect (y-axis) of bat host mass: forearm residual, respectively for *E. dupreanum* and *R. madagascariensis,* on bat fly count. Solid lines (blue for significant effects) correspond to mean effects, with 95% CIs by standard error in translucent shading.

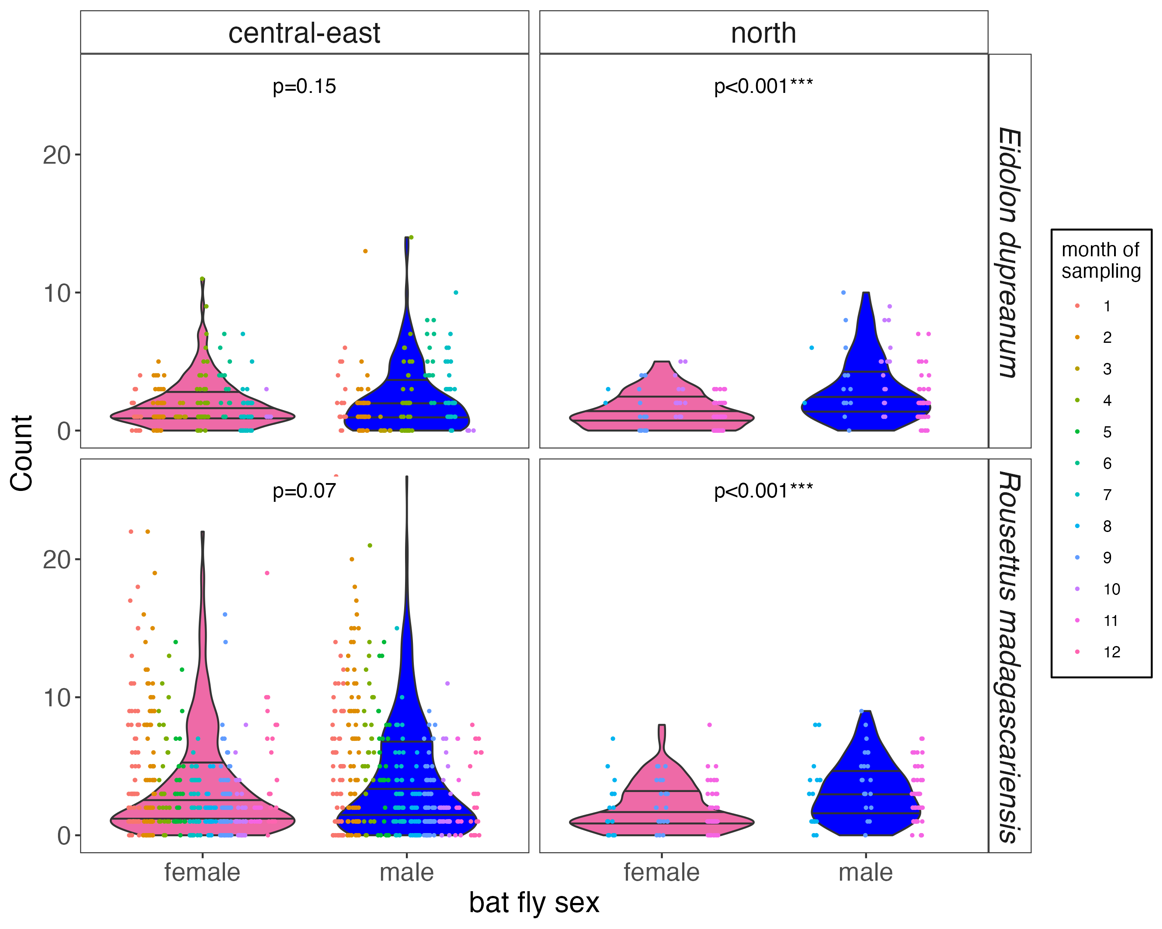

**Fig. S3.** Violin plots showing the range of nycteribiid bat fly abundance, by sex, recovered on *E. dupreanaum* (top) and *R. madgascariensis* (bottom) bat hosts in both central-east and north localities across the range of study dates for the morphological data subset (Feb 2018-Nov 2019). Horizontal lines across each violin show the interquartile range of observed counts, and raw data are plotted as jiggered points onto of the corresponding distribution, colored by month of sampling, according to legend. Reported p-values correspond to a student’s t-test examining the mean abundance of male vs. female nycteribiid count for each bat species host and locality.

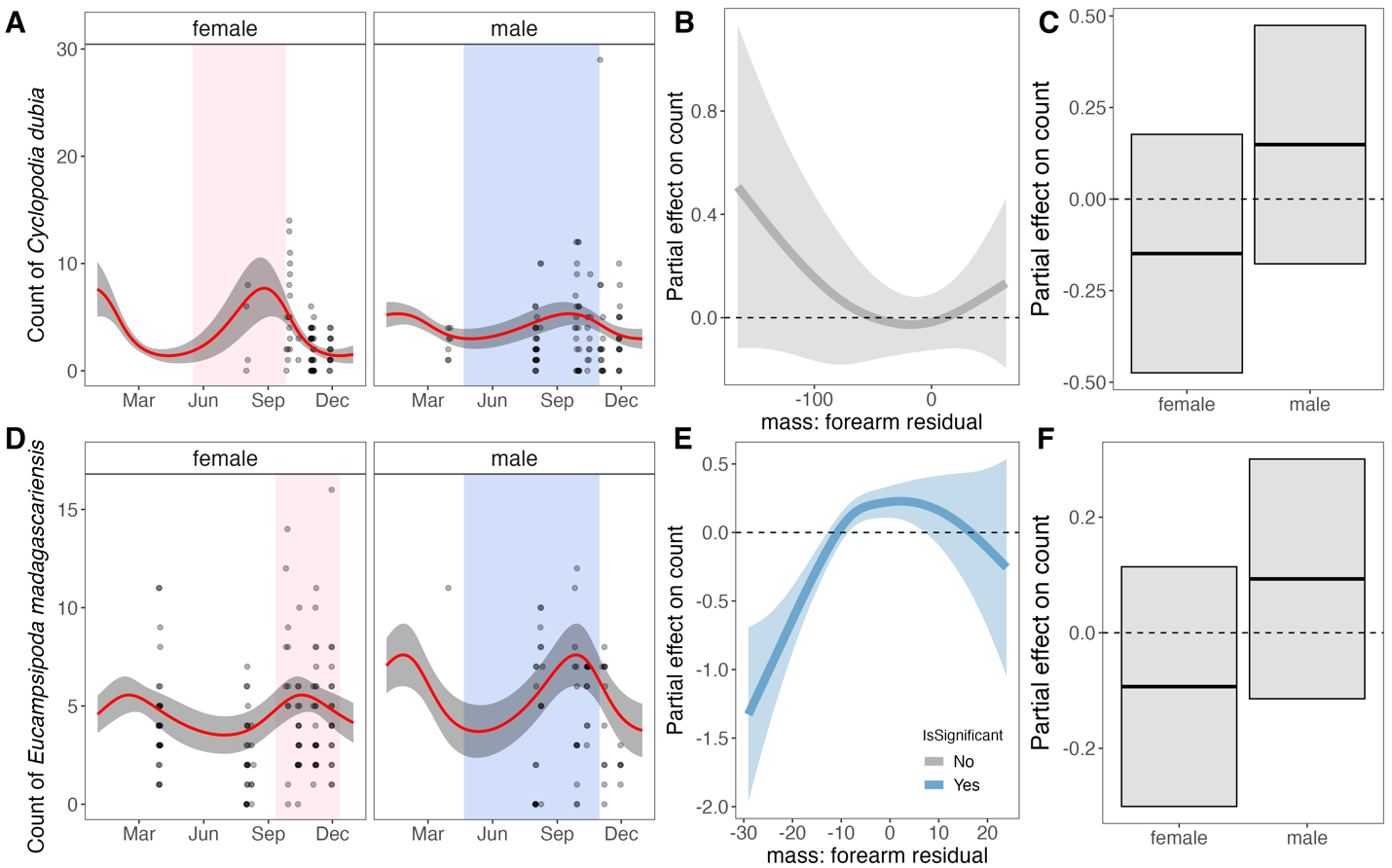

**Fig. S4.** Seasonal variation in the abundance of nycteribiidae bat flies counted on (**A, B**) *E. dupreanum* and (**C,D**) *R. madagascariensis* bats captured at roost sites in northern Madagascar (Ankarana National Park) from field-derived dataset; data were too few for analysis of the morphological data subset from this site. Panels (A) and (D) show seasonal ectoparasite count predictions (red line) from best-fit GAMs for male and female bat hosts of each species, with 95% CI by standard error shaded in gray. Translucent background points correspond to raw data across all years of the data subset (Feb 2018 – Nov 2019), with male bat flies represented in blue and female bat flies in pink. Pink background shading corresponds to the gestation period for each host bat species, adapted from (1) as field data observations indicate a September 26 birth date for *E. dupreanum* in the Ankarana region. Blue background shading corresponds to the nutritionally deficient dry season, which is conserved between central-eastern and northern Madagascar. Panels (B) and (E) show partial effect (y-axis) of bat host mass: forearm residual, respectively for *E. dupreanum* and *R. madagascariensis,* on bat fly count, while panels (C) and (F) show partial effects of host bat sex on bat fly count, again for *E. dupreanum* and *R. madagascariensis*. For panels (B,C,E,F), solid lines (blue for significant effects; gray for non-significant) correspond to mean effects, with 95% CIs by standard error in translucent shading.

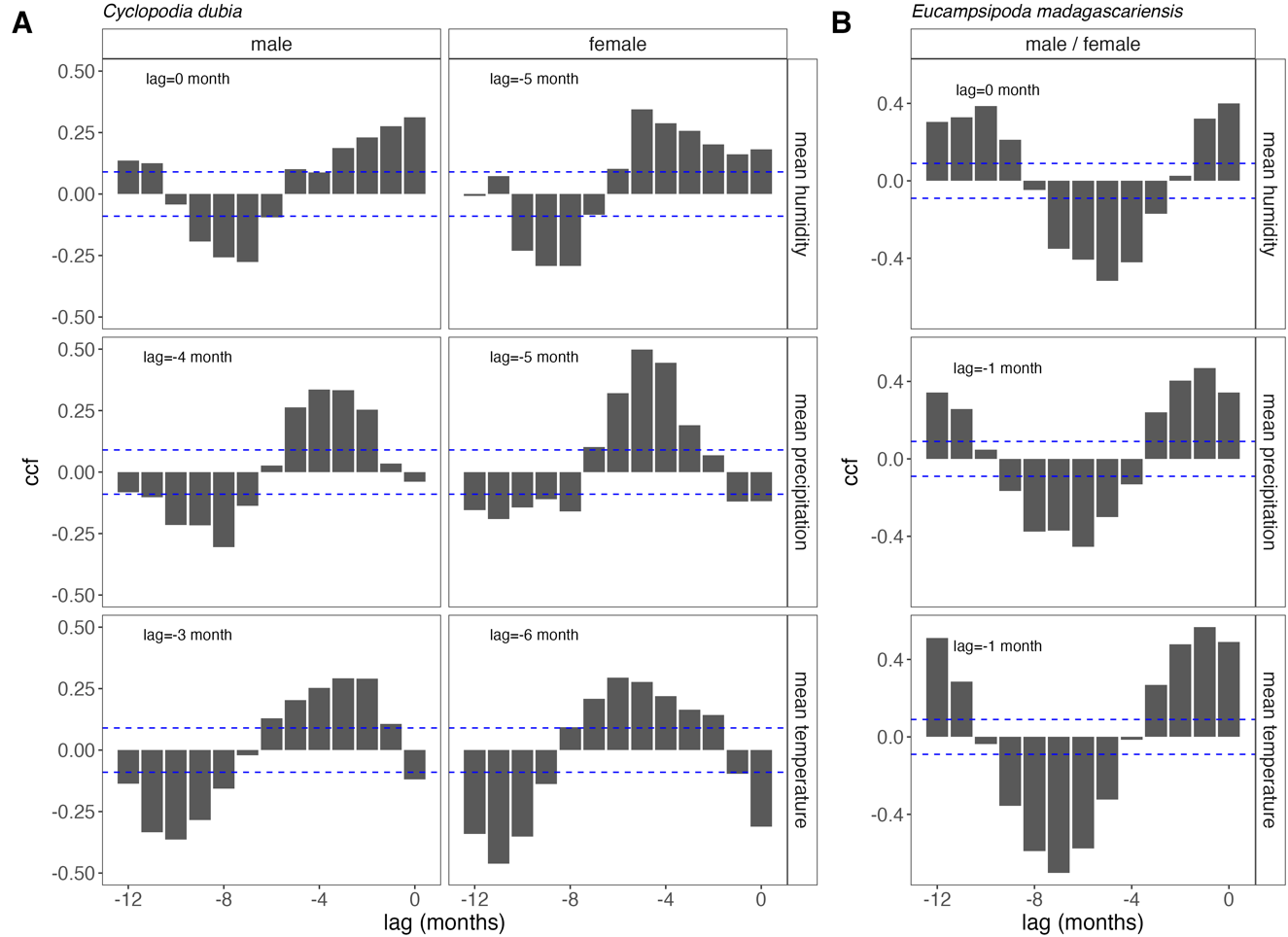

**Fig. S5.** Cross correlation analysis of time series of per bat average monthly nycteribiid bat fly abundance with time series of climate variable (horizontal panels): mean monthy diurnal humidity, mean monthly precipitation rate, mean monthly temperature for (**A**) *C. dubia* parasitizing *E. dupreanum* bats in the Angavokely roost site and (**B**) *E. madagascariensis* bats parasitizing *R. madagascariensis* bats in the Maromizaha roost site. X-axis corresponds to monthly lags up to one year by which climate variables precede ectoparasite burden; while y-axis gives the cross-correlation function (ccf) for maximization. Dashed horizontal lines in blue correspond to thresholds for significant ccf values; the maximal between each pair of time series is listed in the upper lefthand corner of each panel. Separate lags were computed for time series of male and female bat hosts for (A) *C. dubia* on *E. dupreanum.* As maximized lags were equivalent for both male and female bat hosts for (B) *E. madagascariensis* on *R. madagascariensis,* only the lags of the composite dataset are visualized here.

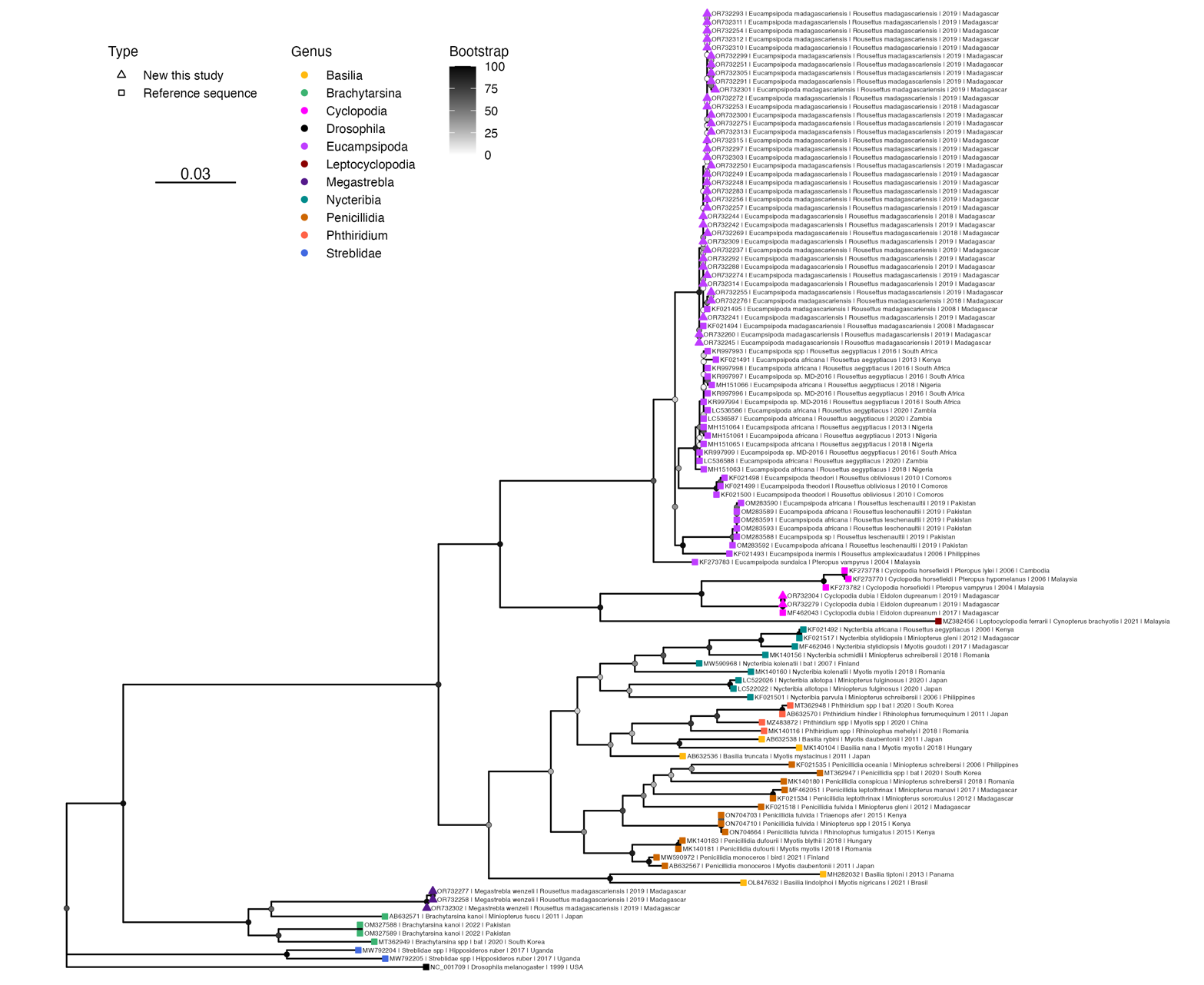

**Fig. S6.** Maximum likelihood phylogeny of COI1 ectoparasite sequences from untrimmed alignment (RAxML-NG, GTR+I+G4) (2). Bootstrap support values computed using Felsenstein’s method (3) are indicated by shaded circles on each node, corresponding to legend. Tip shapes for each sequence are colored by genera, according to the legend, with square shapes corresponding to reference sequences from GenBank and triangle shapes corresponding to new sequences contributed by this study. Tree is rooted in *Drosophila melanogaster,* accession number NC_001709. Branch lengths are scaled by nucleotide substitutions per site, corresponding to the scalebar shown.

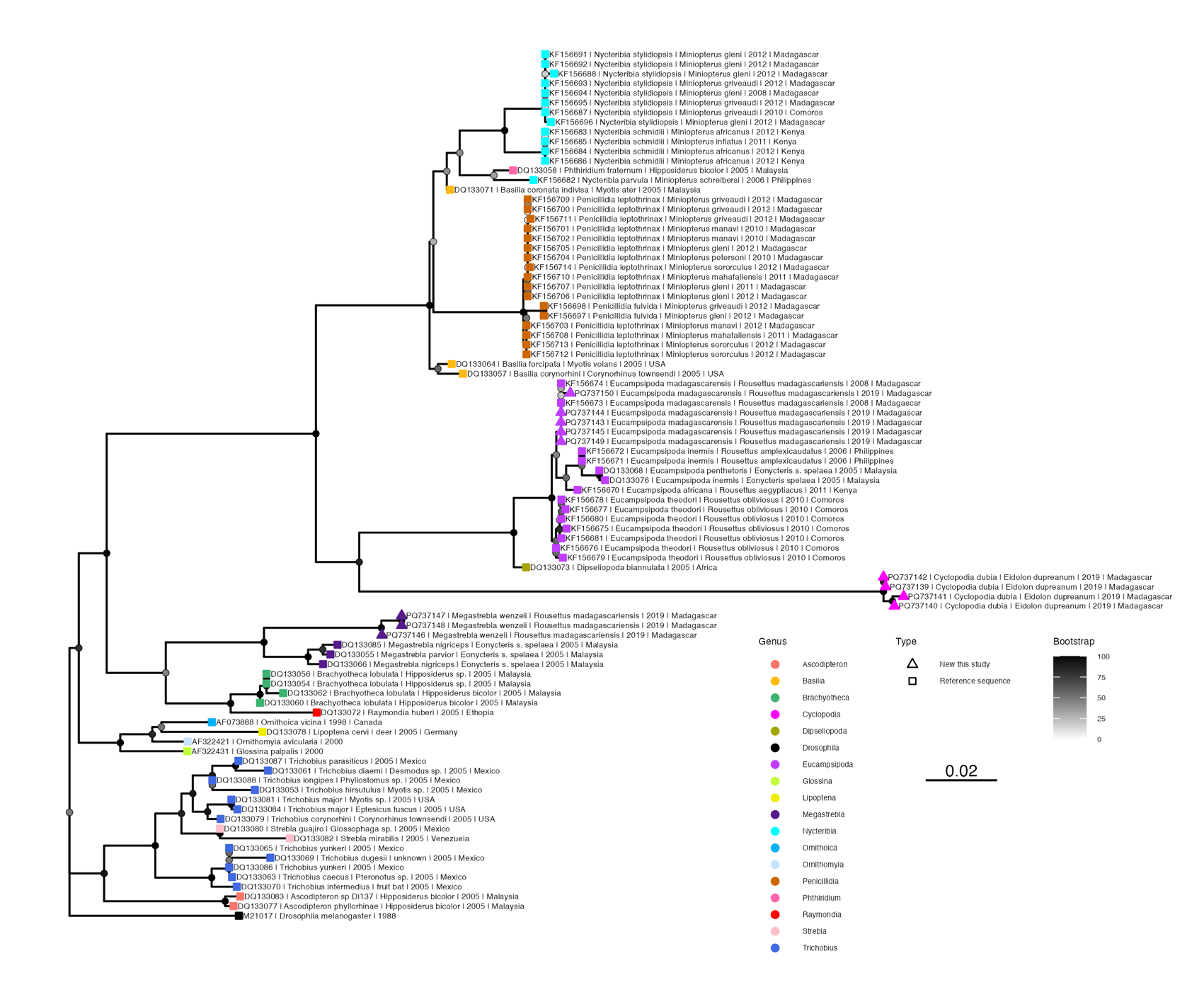

**Fig. S7.** Maximum likelihood phylogeny of 18S ectoparasite sequences from untrimmed alignment (RAxML-NG, TIM2+I+G4) (2). Bootstrap support values computed using Felsenstein’s method (3) are indicated by shaded circles on each node, corresponding to legend. Tip shapes for each sequence are colored by genera, according to the legend, with square shapes corresponding to reference sequences from GenBank and triangle shapes corresponding to new sequences contributed by this study. Tree is rooted in *Drosophila melanogaster,* accession number M21017. Branch lengths are scaled by nucleotide substitutions per site, corresponding to the scalebar shown.
